# Genetic effects of sequence-conserved enhancer-like elements on human complex traits

**DOI:** 10.1101/2022.08.19.504589

**Authors:** Xiang Zhu, Shining Ma, Wing Hung Wong

## Abstract

Non-coding sequences that are evolutionarily conserved and biochemically active offer clues to mechanistic interpretations of human genome-wide association studies (GWAS). However, their genetic effects have not been systematically examined across a wide range of human tissues and traits. Here we develop a simple method to identify functional elements exhibiting high levels of human-mouse sequence conservation and enhancer-like biochemical activity, which scales well to 313 epigenomic datasets across 106 tissues and cell types. Combining these elements with 468 GWAS of European (EUR) and East Asian (EAS) ancestries, we identify tissue-specific enrichments of heritability and causal variants for many traits, as well as candidate genes that are functionally relevant to body mass index (BMI) and schizophrenia but were not reported in previous GWAS. Our findings provide a comprehensive assessment of how sequence-conserved enhancer-like elements affect complex traits, and reinforce the importance of integrating evolutionary and biochemical data to elucidate human disease genetics.

## Introduction

To delineate functional elements ^1^ in the human genome, two major complementary approaches have been developed. The first approach ^2,3^ searches for conserved sequences that remain unchanged over evolution across species (e.g., human and mouse), assuming that mutations therein typically reduce fitness and are thus under negative selection. The second approach ^4,5^ omits the evolutionary conservation and instead identifies sequences of biochemical activity through epigenomic profiling, such as ChIP–seq of H3K27ac to mark active enhancers ^6^. The evolutionary approach has been empowered by the genome sequencing and assembly for a growing number of species ^7,8^. In parallel, genome-wide catalogs of diverse biochemical marks across hundreds of human cell types and tissues have been generated ^9,10^. Collectively, the two approaches have enhanced our understanding of human genome function, especially for the vast non-coding regions that do not encode protein sequences.

Non-coding elements detected by the evolutionary and biochemical approaches are particularly relevant to human disease genetics, as GWAS often implicate non-coding regions ^11^. Non-coding elements with either evolutionary ^12^ or biochemical ^10^ signatures can help prioritize functional variants and yield mechanistic insights at GWAS loci. Further, genomic regions marked by each of the two approaches explain a much larger proportion of complex trait heritability ^13^ than one would expect by the region sizes. Despite the progress, both approaches have limitations to define regulatory elements ^1,6^, and thus each approach alone cannot fully inform regulatory causes of heritable traits.

Recent studies have sought to combine the evolutionary and biochemical approaches to interpret non-coding variation underlying human traits. Integrating evolutionary and biochemical data has proven effective in quantifying fitness consequences for genetic variants ^14,15^, outperforming methods that utilize a single data type ^16^. Besides the totality of phenotypic consequences (fitness), the integrative approach is also useful to elucidate the genetics of a specific trait. DNase I hypersensitivity sites in human fetal brain intersected evolutionarily conserved sequences display a significant excess of de novo mutations exclusively concentrated in neurodevelopmental disorders ^17^. Human orthologues of H3K27ac ^18^ and open chromatin ^19–21^ peaks in mouse brain are enriched for GWAS signals of many brain-related traits in specific brain cell types. H3K27ac peaks in human liver ^22^ show significantly stronger heritability enrichments across 41 complex traits ^23^ when restricted to peaks with sequence age older than the marsupial-placental split ^24^ or peaks with conserved H3K27ac signal across mammals ^22^. These efforts, though promising, only assessed regions with both evolutionary and biochemical signatures for a limited set of tissues and traits. Genetic effects of such regions on a wide range of traits across diverse tissues remain largely unknown, impeding our ability to understand tissue-specific regulation of hereditary traits ^10^.

Here, we develop a human-mouse comparison method to identify human enhancer-like elements that display both sequence conservation and biochemical activity. We apply the method to 313 epigenomic datasets across 106 tissues and cell types, and employ the identified elements to analyze 468 GWAS of EUR and EAS ancestries. These elements not only show strong tissuespecific enrichments of heritability and causal variants for a wide range of traits, but also nominate previously undescribed effector genes for BMI and schizophrenia, revealing additional biological and clinical insights. Overall, we present a simple yet effective strategy to annotate the human genome with complementary lines of evolutionary and biochemical evidence, and demonstrate its utility systematically across a host of tissues and traits.

## Results

### Human-mouse comparisons identify conserved enhancer-like sequences

We developed a simple method to identify active human enhancers that exhibit sequence conservation in the mouse genome (Fig. 1a; Methods). Given a human tissue or cell type (henceforth ‘context’), we first used its H3K27ac profile to empirically determine active enhancers genome wide ^5^, and then intersected them with accessible chromatin regions identified in the same context. Since biochemical activities are not necessarily definitive proof of function ^1,6^, we cautiously termed these regions marked by H3K27ac and chromatin accessibility signals ‘active enhancer-like elements’ (AELEs). Finally, we identified sequence-conserved AELEs by comparing human AELEs with the mouse genome ^25,26^. We specified the sequence conservation level as the minimum proportion of bases mapped to gapless aligned blocks in the mouse genome, with larger values indicating higher conservation levels.

**Fig 1:**
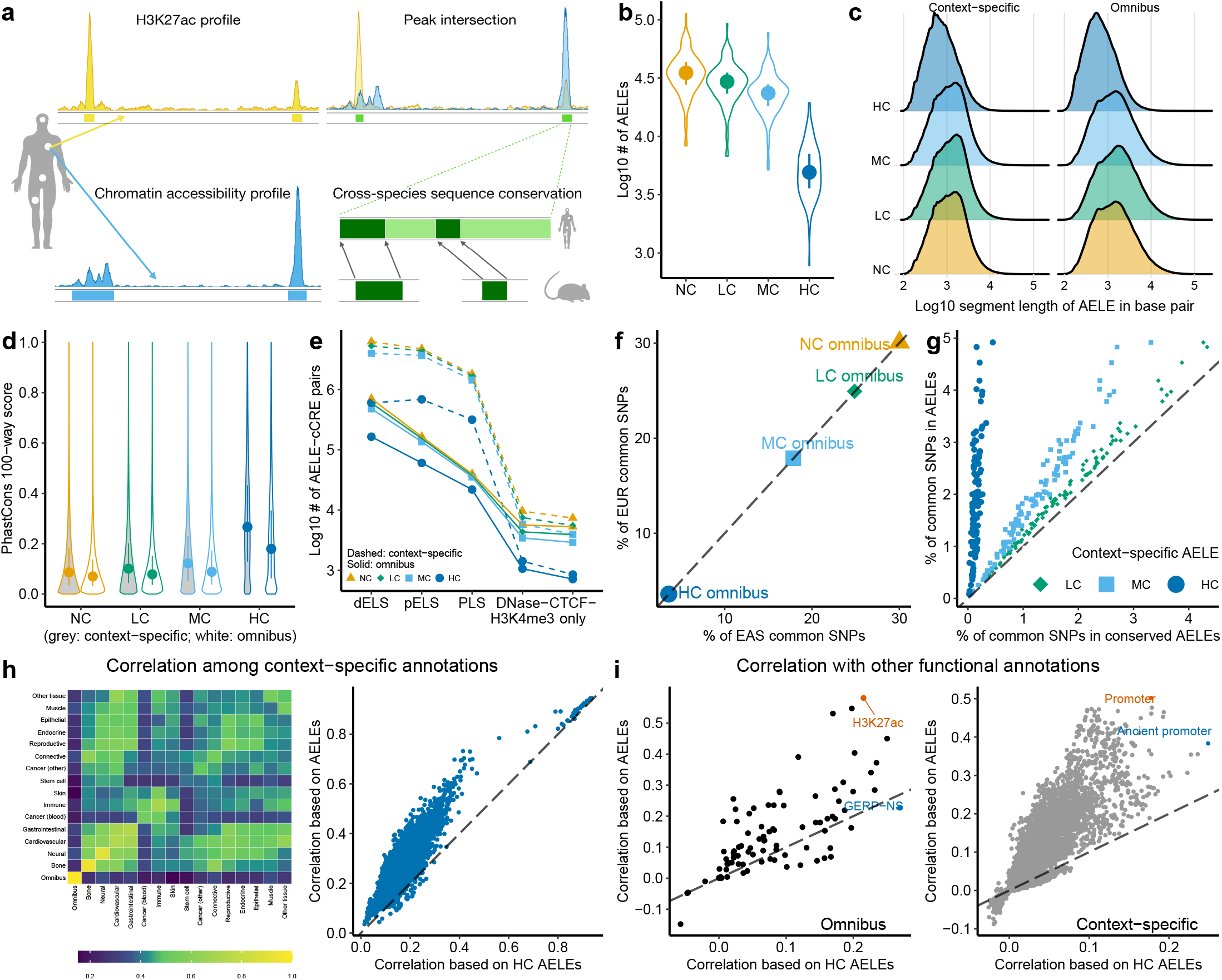
Identify and characterize sequence-conserved enhancer-like elements in the human genome. **a** Schematic of the human–mouse sequence comparison to identify conserved AELEs. **b** Element count distributions for context-specific AELEs without (NC) and with low (LC), moderate (MC) and high (HC) levels of sequence conservation across 105 contexts. **c** Segment length distributions for NC, LC, MC and HC AELEs. **d** Evolutionary conservation score distributions for NC, LC, MC and HC AELEs. **e** Numbers of overlapping pairs of AELEs and ENCODE cCREs, stratified by sequence conservation levels and cCRE classifications (dELS & pELS: distal & proximal enhancer-like signature; PLS: promoter-like signature). **f** Percentages of EUR (*y*-axis) and EAS (*x*-axis) common SNPs that fall inside NC, LC, MC and HC omnibus AELEs. **g** Given a context, percentages of EUR common SNPs that fall inside context-specific AELEs without (*y*-axis) and with (*x*-axis) sequence conservation. **h** The heatmap shows the maximum pairwise correlations of common SNP annotations based on NC context-specific AELEs from two context groups (Table S1). The group labeled as ‘other tissues’ consists of kidney, liver and lung. The scatter plot shows the pairwise correlations of common SNP annotations based on NC (*y*-axis) and HC (*x*-axis) context-specific AELEs, where each point denotes a pair of contexts. **i** Each point denotes a pair of an AELE annotation and one of the 96 known annotations, with *y* and *x*-axis indicating correlations between the known SNP annotation and the SNP annotation based on NC and HC AELEs respectively. To simplify visualization, only points with the largest *y* and *x*-axis values are labeled by the corresponding known annotations in each panel. For **b** and **d**, each point denotes a median and each line denotes an interquartile range. For **f**-**i**, dashed lines have intercept 0 and slope 1.

We applied this method to context-matched H3K27ac and chromatin accessibility profiles across 106 contexts in humans (Table S1). We first generated AELEs for each context. The median AELE count per context is 35065 (range: 8309-112362). Across all AELEs, the median segment length is 1119 bp (range: 100-194631 bp). We then created three sets of conserved AELEs in the same context by setting the sequence conservation level as 0.1, 0.5 and 0.9, denoted as lowly (LC), moderately (MC) and highly-conserved (HC) AELEs, respectively. As expected, there are fewer HC AELEs than AELEs in the same context (median decrease: 29296; Fig. 1b) and HC AELEs are shorter than AELEs (median decrease: 450 bp; Fig. 1c).

For each conservation level (none, LC, MC, HC), we further aggregated all context-specific AELEs across 106 contexts and merged overlapping segments into non-overlapping ones to produce an “omnibus” version (Fig. S1a), resulting in 338743 unique omnibus human AELEs as well as 247504 LC, 222160 MC and 132717 HC omnibus AELEs (Fig. S1b). The increased element counts are expected, because omnibus AELEs accumulate AELEs from diverse contexts, most of which are specific to a single context. Specifically, 52.1% of HC omnibus AELEs are indeed HC AELEs from one context (Fig. S1c) and 57.6% consist of HC AELEs from one context group (Fig. S1d).

To assess their evolutionary and biochemical relevance, we overlapped AELEs with the 100-vertebrate phastCons ^16^ scores and the ENCODE ^9^ human candidate cis-regulatory elements (cCREs). Reassuringly, HC omnibus AELEs are more evolutionarily conserved than omnibus AELEs (median increase: 0.11; one-sided Wilcoxon *P* < 2.2 × 10^−308^; Fig. 1d). More than 90% of the omnibus AELE-cCRE pairs contain enhancer-like signatures irrespective of sequence conservation (Fig. 1e). We observed similar patterns for contextspecific AELEs. Together, the results show that AELEs are often biochemically active, while HC AELEs display evolutionary and biochemical signals.

To assess their regulatory functions, we tested motif enrichments in AELEs (Methods). Despite their smaller coverage of the human genome compared with AELEs (Fig. 1b-c), conserved AELEs contain many significantly enriched motifs (Table S2). Specifically, we identified 151 unique motifs in HC AELEs showing strong enrichments (≥ 2-fold and *P* ≤ 1.0 × 10^−12^) in at least one context against the GC-matched random background. To avoid confounding caused by context specificity, we repeated this analysis with the background being AELEs in the same context and identified enrichments for 186 unique motifs. Some enriched motifs are relevant to the context from which HC AELEs are derived. For example, HC AELEs derived from neural progenitors are enriched for a DBX2 motif (2.09-fold, *P* = 1.0 × 10^−100^), consistent with the brain-specific mRNA expression of DBX2 and its regulatory role in age-related neurogenic decline ^27^. The motif enrichments confirm high concentrations of regulatory sequences in conserved AELEs, forming a basis to interpret non-coding variation.

To capture their genetic variation, we mapped AELEs to biallelic autosomal single-nucleotide polymorphisms (SNPs) with minor allele frequency (MAF) ≥ 0.05 (henceforth ‘common SNPs’) in EUR and EAS ancestries (Methods). In total, 30.1% of 5961159 EUR common SNPs lie within omnibus AELEs, 24.9% within LC, 17.9% within MC and 3.6% within HC omnibus AELEs. These percentages are the same up to three decimal places for 5469053 EAS common SNPs (Fig. 1f). We also mapped context-specific AELEs to the common SNPs and observed a similar trend of AELEs with a higher level of sequence conservation covering fewer common SNPs (Fig. 1g), consistent with the patterns of element count (Fig. 1b) and segment length (Fig. 1c).

To investigate context specificity, we examined correlations between AELE annotations of common SNPs for all context pairs (Fig. 1h; Table S3). Reassuringly, correlations are generally stronger (average increase: 0.12; onesided Wilcoxon *P* = 4.4 × 10^−35^) when AELEs belong to the same context group (e.g., ascending aorta and tibial artery, Cramér’s *V* = 0.73) than when they are in distant groups (e.g., neural progenitor and HCASMC, *V* = 0.08). The correlations are high in different but related contexts, such as immune and blood cancer groups (both rich in immune cells, *V* = 0.53), cardiovascular and gastrointestinal groups (both rich in muscle and connective cells, *V* = 0.69). Conserved AELEs produce concordant results for the same context pairs (Pearson’s *R* = 0.87 − 0.99; Fig. 1h), showing that conserved AELEs preserve the context specificity of AELEs.

We further correlated AELE annotations with 96 functional annotations ^23^ of the common SNPs (Fig. 1i; Table S4). The omnibus AELE annotation is weakly correlated with most of the 96 annotations (median *V* = 9.9 × 10^−2^; ^13^ ×10^−4^ ≤ *V* ≤ 0.58). The strongest correlation is for a context-merged H3K27ac annotation, consistent with the construction of omnibus AELEs (Fig. 1a; Fig. S1a). Compared to the omnibus AELE annotation, the HC omnibus AELE ^28^ annotation is far less correlated with existing annotations (median *V* = 6.5 × 10^−2^; 9.0 × 10^−5^ ≤ *V* ≤ 0.27; one-sided Wilcoxon *P* = 2.3 × 10^−9^). The strongest correlation is for an evolutionary constraint annotation, consistent with the role of sequence conservation in identifying HC AELEs. We observed even weaker correlations between context-specific AELE annotations and existing annotations (AELEs: 1.1×10^−7^ ≤ *V* ≤ 0.50; HC AELEs: 1.3×10^−6^ ≤ *V* ≤ 0.25). Overall, SNP annotations based on AELEs, especially HC AELEs, are not strongly correlated with existing annotations, suggesting their potential to capture additional genetic signals of complex traits.

### HC AELEs explain heritability independent of known annotations

Enrichments of common SNP heritability for complex traits have been shown in many genomic annotations ^13,23^. Because of their potential functions (Fig. 1d-e) and weak correlations with existing annotations (Fig. 1i), we hypothesized that omnibus AELEs could be enriched in heritability, independent of contributions from known annotations. To test this hypothesis, we used stratified LD score regression ^13^ (S-LDSC) to analyze 468 GWAS (Table S5) and 4 SNP annotations of omnibus AELEs (Methods; Table S6). For each GWAS, we first applied S-LDSC to the annotation of omnibus AELEs while conditioning on 96 functional annotations ^23^. We then analyzed each of three annotations of conserved (LC, MC, HC) omnibus AELEs with S-LDSC while conditioning on the omnibus AELEs and 96 previous annotations. For each GWAS and annotation, we summarized the S-LDSC analysis by: (1) heritability enrichment, defined as the proportion of heritability explained by SNPs in the annotation divided by the proportion of SNPs in the annotation; (2) standardized effect size (*τ*^⋆^), defined as the proportionate change in per-SNP heritability associated with a 1-standard deviation increase of the annotation, conditioned on all other annotations. Given an annotation, a significant heritability enrichment > 1 indicates the marginal effect, whereas a significant *τ*^⋆^ > 0 indicates the joint effect unique to the annotation.

In the meta-analysis across all 468 GWAS, we observed a significant heritability enrichment for omnibus AELEs (1.71-fold, one-sided *P =* 6.9 × 10^−263^), consistent with previous work ^13,23^. We also found omnibus AELEs uniquely informative for per-SNP heritability conditional on 96 known annotations, as quantified by *τ*^⋆^ = 0.029 (one-sided *P* = 1.7 × 10^−2^). When analyzing conserved omnibus AELEs, we identified a rising signal strength along the sequence conservation level (Fig. 2a). Specifically, we obtained 4.88-fold heritability enrichment (*P* = 5.4 × 10^−325^) for HC omnibus AELEs, compared to 2.29-fold (*P* = 3.0 × 10^−277^) for MC and 1.93-fold (*P* = 1.7 × 10^−341^) for LC. Conditional on omnibus AELEs and 96 functional annotations, we estimated *τ*^⋆^ = 0.255 (*P* = 4.8 × 10^−65^) for HC omnibus AELEs, compared to *τ*^⋆^ = 0.101 (*P* = 2.8 × 10^−13^) for MC and *τ*^⋆^ = 0.086 (*P* = 1.8 × 10^−6^) for LC. Together the results demonstrate a significant effect of HC omnibus AELEs on heritability, which is not explained by omnibus AELEs or known annotations.

**Fig 2:**
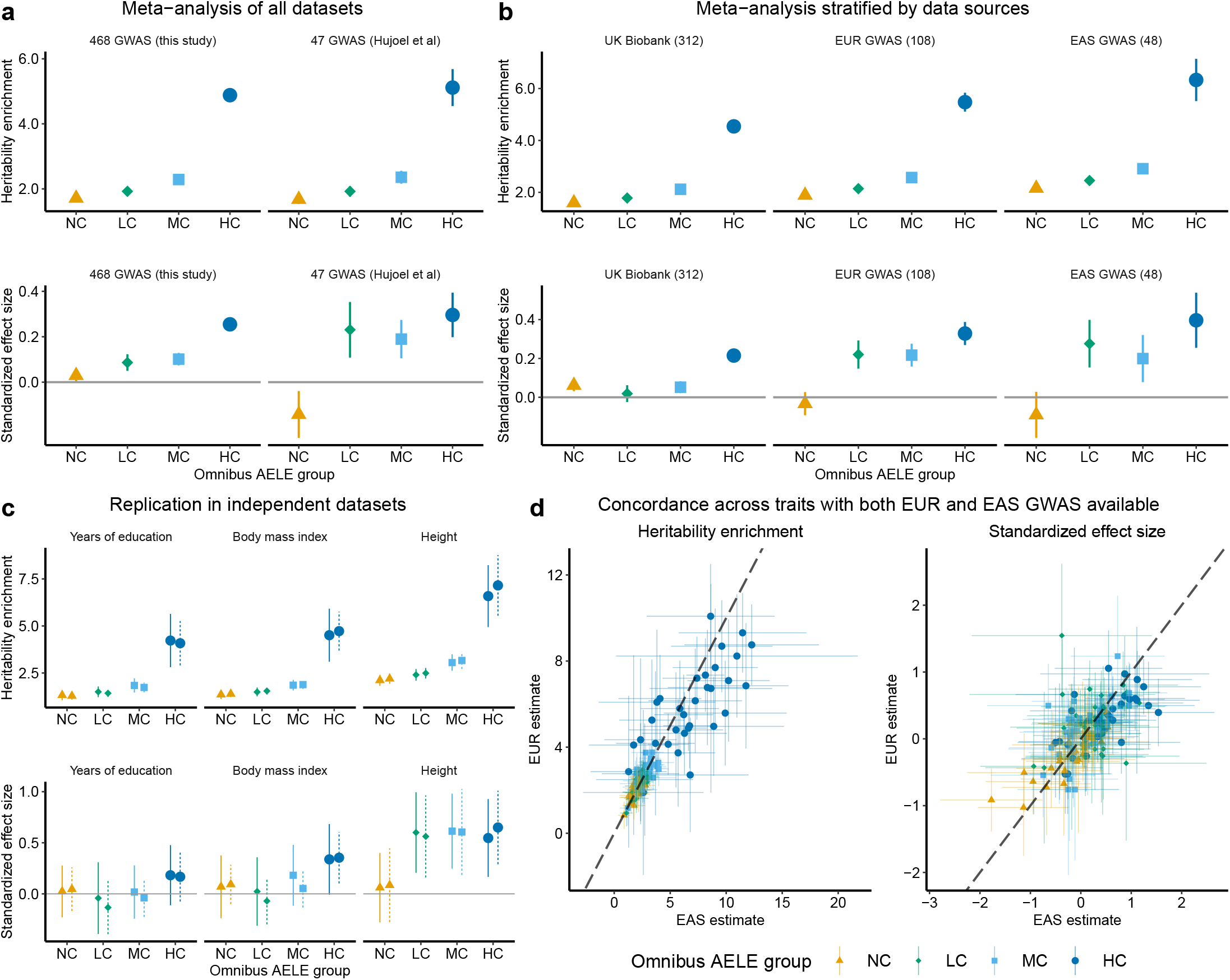
Quantify common SNP heritability enrichments for omnibus AELEs with varying levels of sequence conservation. We show heritability enrichment and standardized effect size (*τ*^∗^) estimates for 4 SNP annotations based on the omnibus AELEs without (NC) and with sequence conservation (LC, MC, HC). The estimates for omnibus AELEs (NC) are conditional on 96 known annotations. The estimates for LC, MC, HC omnibus AELEs are conditional on omnibus AELEs and 96 known annotations. **a** Estimates meta-analyzed across 468 GWAS datasets curated for this study and a previously defined set of 47 independent GWAS datasets respectively. **b** Estimates meta-analyzed across 312 UK Biobank, 108 EUR and 48 EAS GWAS datasets respectively. Numerical results for **a** and **b** are available in Table S7. **c** Comparison of heritability enrichment and *τ*^∗^ estimates based on 2 independent EUR GWAS for 3 representative traits. **d** Heritability enrichment and *τ*^∗^ estimates for 4 omnibus AELE annotations across 34 traits that each has EUR and EAS GWAS available. Numerical results for **c** and **d** are available in Table S6. For **a**-**d**, each point denotes an estimate and each error bar denotes ±1.96 SE. For **c**-**d**, dashed lines have intercept 0 and slope 1.

As a sensitivity analysis, we restricted the meta-analysis to a previously described set ^23^ of 47 independent datasets (Fig. 2a). Despite larger standard errors (SEs) caused by the fewer datasets meta-analyzed, we obtained similar heritability enrichments for all four annotations (*P* ≥ 0.44 for difference). We also estimated similar *τ*^⋆^ for HC omnibus AELEs (*P* = 0.43 for difference), highlighting the robustness of results based on this annotation.

We further meta-analyzed results (Table S7) stratified by study populations (Fig. 2b) and trait categories (Fig. S2), reaching two conclusions in concordance with the full analysis (Fig. 2a). First, HC omnibus AELEs have stronger heritability enrichments than omnibus AELEs (2.12 − 6.13 fold increase). Second, HC omnibus AELEs have significantly positive effect sizes (median *τ*^⋆^ = 0.33 and *P* = 3.6 × 10^−5^ across 28 strata) conditional on omnibus AELEs and 96 known annotations. Across three populations, the metaanalysis of 48 EAS GWAS produced the strongest enrichment for HC omnibus AELEs (6.33-fold, *P* = 1.6 × 10^−27^; *τ*^⋆^ = 0.396, *P* = 2.2 × 10^−8^). Within EUR GWAS, the meta-analysis of 8 cardiovascular traits produced the strongest enrichment (8.13-fold, *P* = 6.2 × 10^−6^; *τ*^⋆^ = 0.682, *P* = 6.8 × 10^−8^). Within UK Biobank, the meta-analysis of 19 medication use traits produced the strongest enrichment (6.04-fold, *P* = 6.6 × 10^−11^; *τ*^⋆^ = 0.525, *P* = 2.2 × 10^−13^). Despite quantitative differences the qualitative finding remains the same: HC AELEs are more informative than AELEs for heritability enrichment.

To assess replicability, we examined S-LDSC results of 13 traits that each had two independent EUR GWAS with comparable sample sizes. For all four annotations of omnibus AELEs, we obtained similar results between two independent datasets of the same trait (*P* > 0.05/13 for difference; Fig. 2c).

To evaluate transferability of our findings across populations, we compared results of 34 traits that each had EUR and EAS GWAS available. Across annotations and traits, we obtained concordant estimates between EUR and EAS (Fig. 2d; heritability enrichment: *R* = 0.91, *P* = 7.6 × 10^−52^; *τ*^⋆^: *R* = 0.63, *P* = 1.9 × 10^−16^). Further, we found no evidence of population heterogeneity for the same annotation and trait (*P* > 0.05/34 for difference). The estimates tend to be smaller in EUR than EAS (heritability enrichment: slope = 0.93, SE = 0.052; *τ*^⋆^: slope = 0.86, SE = 0.068; Methods), which is consistent with our meta-analysis stratified by populations (Fig. 2b) as well as a recent EAS-EUR comparison across 29 traits and 100 regulatory annotations ^29^. Overall, the results indicate comparable heritability enrichments for EUR and EAS in omnibus AELEs, and consistently stronger enrichments in HC omnibus AELEs than in omnibus AELEs for both populations.

### HC AELEs show context-specific heritability enrichments

Having established the strong heritability enrichment for HC omnibus AELEs, we next assessed context-dependent enrichments for HC AELEs (Table S8). Specifically, we analyzed the annotation of HC context-specific AELEs from each context against each GWAS with S-LDSC, while conditioning on AELEs in the same context and 96 previous annotations (Methods). We quantified the significance of context-specific enrichment by a one-sided *P*-value that tests *τ*^⋆^ > 0, controlling for effects of all other annotations.

We first meta-analyzed results across groups of related traits and contexts. For many trait groups, we observed top-ranked enrichments in HC AELEs derived from contexts highly relevant to the traits (Fig. 3a; Fig. S3; Table S9). HC AELEs derived from the nervous system show strong enrichments for mental disorders (*τ*^⋆^ = 0.556, *P* = 1.7 × 10^−59^) and a wide range of traits related to behavior (*τ*^⋆^ = 0.451, *P* = 1.4 × 10^−125^), sleep (*τ*^⋆^ = 0.422, *P* = 2.5 × 10^−76^), reproduction (*τ*^⋆^ = 0.367, *P* = 1.1 × 10^−8^) and diet (*τ*^⋆^ = 0.359, *P* = 1.3 × 10^−162^). HC AELEs derived from the immune system show strong enrichments for immune diseases (*τ*^⋆^ = 0.543, *P* = 9.4 × 10^−24^) and blood cell traits (*τ*^⋆^ = 0.226, *P* = 1.0 × 10^−9^). Other examples include bone for bone traits (*τ*^⋆^ = 0.332, *P* = 1.1 × 10^−29^), connective tissue for early growth traits (*τ*^⋆^ = 0.456, *P* = 7.2 × 10^−11^), and kidney for kidney traits (*τ*^⋆^ = 0.998, *P* = 4.2 × 10^−9^). The significantly positive *τ*^⋆^ estimates indicate that HC AE-LEs provide additional information about heritability conditional on AELEs in the same context. Further, top enrichments of HC context-specific AELEs are consistently stronger than enrichments of HC omnibus AELEs based on the same GWAS, recapitulating the tissue selectivity of heritable traits ^10,13^.

**Fig 3:**
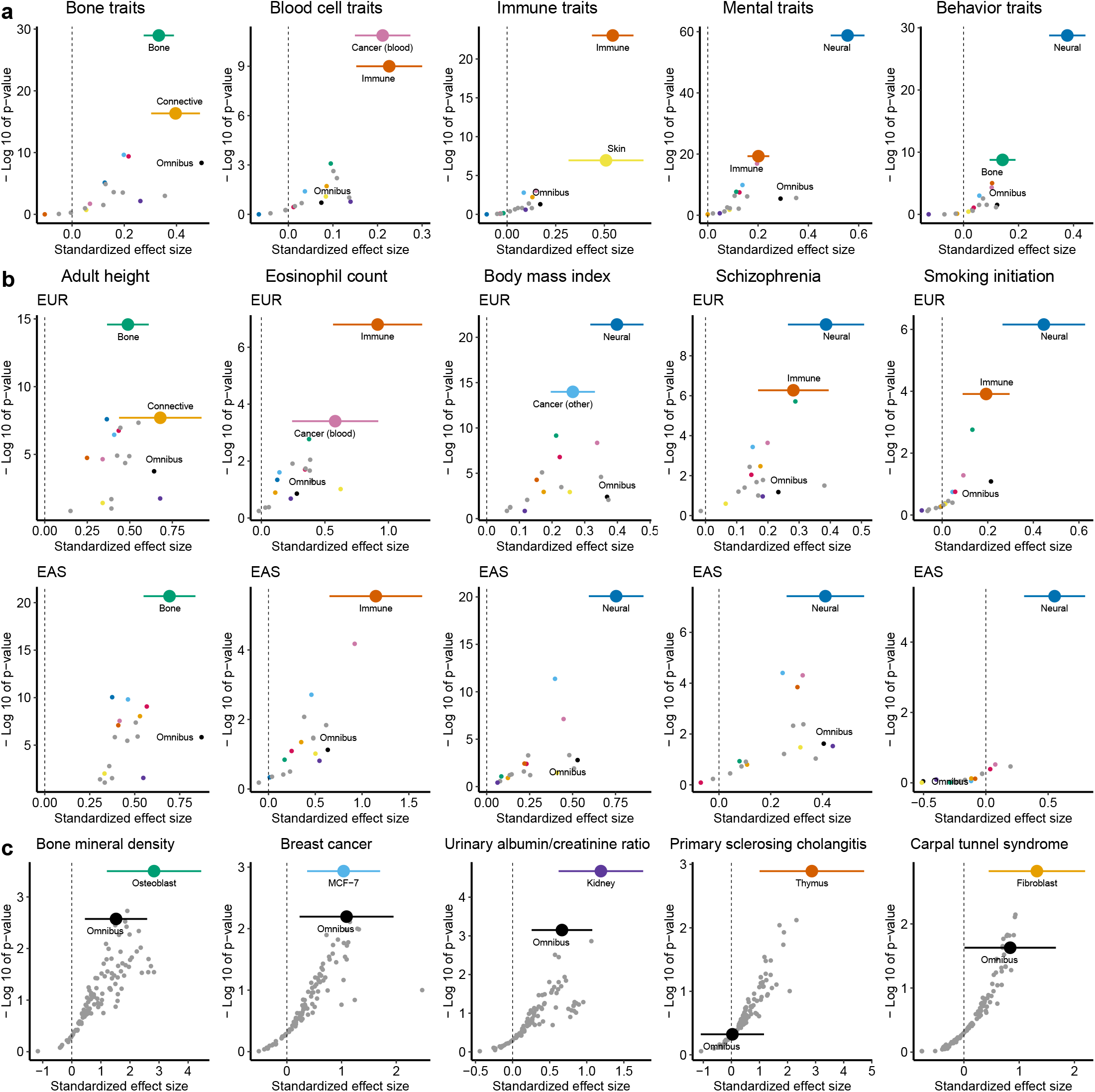
Assess common SNP heritability enrichments for HC context-specific AELEs. For HC context-specific AELEs from each of the 105 contexts, we compute the standardized effect size (*τ*^⋆^) estimate and *P*-value for testing *τ*^⋆^ > 0 conditional on AELEs in the same context and 96 known annotations. **a** Estimates meta-analyzed within each of the 17 context groups (Table S1) for 5 groups of related traits in EUR GWAS. Additional results are shown in Fig. S3. **b** Estimates meta-analyzed within each of the 17 context groups for 5 traits that have both EUR and EAS GWAS available. Numerical results for **a** and **b** are available in Table S9. **c** Estimates of 105 individual contexts for 5 exemplary traits. Numerical results are available in Table S8. For **a**-**c**, each point denotes an estimate and each error bar denotes ±1.96 SE. The color legend is provided in Fig. S1i.

We also observed context-dependent enrichments in the meta-analysis of related contexts for a single trait. Specifically, for 34 traits with both EUR and EAS GWAS available, we often identified the strongest enrichments of HC AELEs from the same context group (Fig. 3b). For example, HC AELEs derived from the nervous system show the strongest enrichments in EUR and EAS for BMI, schizophrenia and smoking initiation. Other examples are bone for adult height and immune system for eosinophil count. Across all contexts and traits, we obtained concordant estimates of *τ*^⋆^ between EUR and EAS (*R* = 0.53, *P* = 1.4×10^−253^) and found no evidence of population heterogeneity (*P* > 0.05/(34 × 105) for difference). As in the omnibus results (Fig. 2d), we estimated smaller *τ*^⋆^ for HC context-specific AELEs in EUR than in EAS (slope = 0.69, SE = 0.018). Overall, the results showcase the transferability of heritability enrichments for HC context-specific AELEs across populations.

Lastly, we examined individual contexts for a given trait (Fig. 3c). As expected, individual enrichments are weaker than meta-analyzed enrichments, but the top-ranked ones still inform trait-relevant contexts. Some top-ranked enrichments are straightforward to interpret, such as kidney for urine albumin-creatinine ratio, osteoblast for bone mineral density and MCF-7 for breast cancer. Some top-ranked enrichments are less direct but functionally relevant nonetheless. HC AELEs derived from thymus (where T cells mature) show the strongest enrichment among 105 contexts for primary sclerosing cholangitis (PSC), followed by HC AELEs derived from T cells. This adds to the considerable evidence linking T cells to the pathogenesis of PSC ^30^. HC AELEs derived from fibroblast (connective tissue) show the strongest enrichment for carpal tunnel syndrome (CTS), consistent with the fibrosis of subsynovial connective tissue in CTS patients ^31^.

### HC AELEs capture more heritability than H3K27ac-conserved AELEs

Our identification of HC AELEs differs from previous work of detecting human enhancers aligned with H3K27ac signals in other mammals ^22^. To compare the two approaches, we exploited 14 contexts that each had data of human chromatin accessibility, human and mouse H3K27ac available (Table S10). For each context, we created two sets of HC AELEs based on conservation with mouse genome sequences (Fig. 1a) and H3K27ac peaks (Fig. S4a; Methods) respectively, and then assessed heritability enrichments on the same GWAS (Table S11). Though using less information, we observed stronger heritability enrichments for HC AELEs based on the mouse genome (10.9-fold, *P* = 5.4 × 10^−14^; *τ*^⋆^ = 0.158, *P* = 8.9 × 10^−215^) than for HC AE-LEs based on mouse H3K27ac peaks (8.4-fold, *P* = 4.9 × 10^−15^; *τ*^⋆^ = 0.079, *P* = 6.1×10^−121^) in the meta-analysis across GWAS and contexts (Table S12). We obtained similar results when restricting to an independent set of GWAS (Fig. S4b) or individual contexts (Fig. S4c). Besides capturing more heritability than H3K27ac-conserved AELEs, HC AELEs does not need human-mouse H3K27ac data in the same context, thus improving applicability.

### HC AELEs harbor an excess of likely causal variants

Besides heritability enrichment, we examined fine-mapped GWAS variants in AELEs. Specifically, we intersected omnibus AELEs and conserved counterparts (LC, MC, HC) with 515848 fine-mapped variants ^32^ of 94 traits whose posterior inclusion probabilities (PIPs) were estimated by FINEMAP and SUSIE independent of any functional annotation (Methods). For each trait and annotation, we computed the fraction of fine-mapped variants residing in the elements with PIPs above a given threshold. We further compared these fractions with the fraction of all fine-mapped variants that had PIPs above the same threshold in the same trait, to quantify enrichments.

Across 94 traits (Fig. 4a), we observed consistently larger fractions (median increase: 0.5% −9.2%) and stronger enrichments (median increase: 5.6× 10^−3^ − 1.7) of fine-mapped variants in HC omnibus AELEs than in omnibus AELEs, as PIP thresholds varied from 0 to 0.5. For example, across 94 traits we obtained a median 13.9% of variants with SUSIE-estimated PIP ≥ 0.1 among the fine-mapped variants residing in HC omnibus AELEs, compared to 10.1% for MC, 9.4% for LC, 8.9% for omnibus AELEs and 6.1% for the whole genome. We obtained highly concordant results between SUSIE and FINEMAP (e.g., *R* = 0.99 for HC omnibus AELEs), confirming the robustness of our findings to fine-mapping methods. We also observed similar patterns in individual traits (Fig. 4b; Table S13). Altogether, the results demonstrate a significant enrichment of putative causal variants in HC omnibus AELEs.

**Fig 4:**
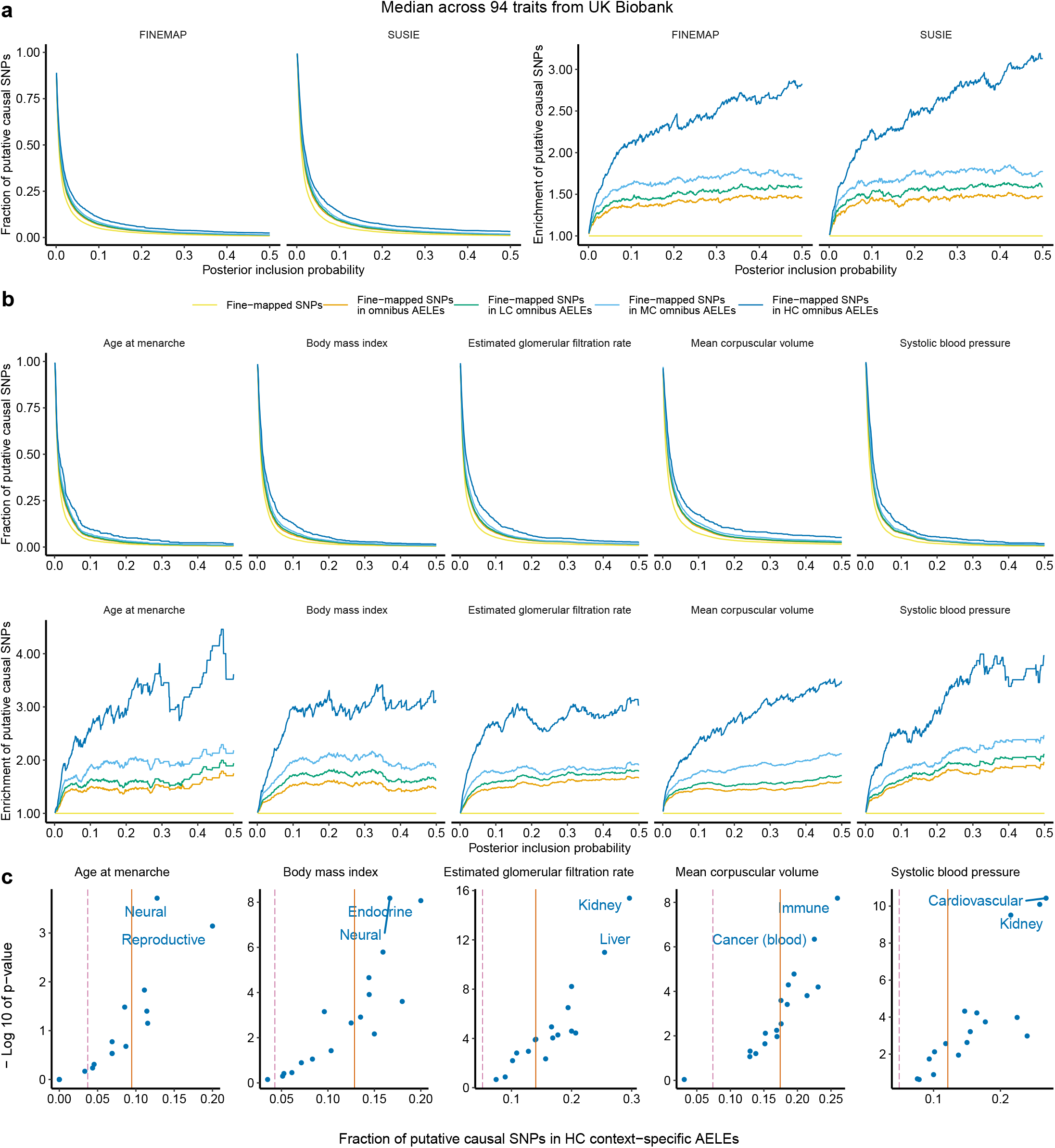
Overlap fine-mapped GWAS variants with AELEs. For each of the 94 traits, we compute fractions of putative causal SNPs that fall inside in the omnibus AELEs without and with sequence conservation, and then compare them against the fraction of putative causal SNPs among all fine-mapped SNPs for the trait to assess enrichments. A fine-mapped SNP is “putative causal” for a trait if this SNP has PIP above a given threshold. **a** Median fractions and enrichments across 94 traits for 2 fine-mapping methods. Numerical results are available in Table S13. **b** Individual fractions and enrichments for 5 traits based on SUSIE. Additional results are available in Table S13. **c** Fractions of putative causal SNPs (SUSIE PIP≥ 0.1) that fall inside HC context-specific AELEs from each of the 17 context groups for 5 traits. The solid and dashed lines denote the fractions of putative causal SNPs for a given trait that fall inside HC omnibus AELEs and the whole genome respectively. Additional results are available in Table S14.

To characterize the context specificity of fine-mapping enrichments, we intersected HC AELEs of 17 context groups (Table S1) with the 515848 finemapped variants (Fig. 4c; Table S14). For each trait and group, we calculated the fraction of fine-mapped variants residing in HC AELEs from this context group with PIPs above 0.1. We compared this fraction with the fraction of all fine-mapped variants that had PIPs above 0.1 in the same trait, producing a one-sided bionomial *P*-value to quantify enrichments. Similar to heritability enrichments (Fig. 3), context-specific enrichments of fine-mapped variants in HC AELEs highlight trait-relevant contexts. Fine-mapped variants for BMI show a stronger enrichment in HC AELEs from the neural (16.7%, *P* = 6.6 × 10^−9^) and endocrine (20.0%, *P* = 8.5 × 10^−9^) groups than the HC omnibus AELEs (12.9%). For estimated glomerular filtration rate, fine-mapped variants are strongly enriched in HC AELEs from the kidney group (29.6%, *P* = 4.1×10^−16^; omnibus: 14.0%), Other examples include HC immune-related AELEs for blood cell phenotypes (18.8−45.0%, *P* = 6.5×10^−9^ −7.6×10^−6^; omnibus: 13.6−25.2%) and HC cardiovascular-related AELEs for blood pressure traits (17.1 − 26.8%, *P* = 3.7 × 10^−11^ − 8.6 × 10^−7^; omnibus: 10.8 − 12.6%).

### HC AELEs aid prioritization of trait-associated regulatory elements

To prioritize trait-associated regulatory elements based on conserved AELEs, we extended RSS-NET ^33^, a method that simultaneously infers genetic enrichments and associations from GWAS summary statistics and genomic annotations (Methods). We applied this extension to analyze omnibus AELEs in GWAS of BMI ^34^ and schizophrenia ^35^. As a sanity check, we examined enrichments produced by RSS-NET in each trait (Fig. 5a; Fig. 6a). Reassuringly, HC omnibus AELEs show stronger enrichments of genetic associations than omnibus AELEs for both traits, mirroring the pattern of S-LDSC results.

**Fig 5:**
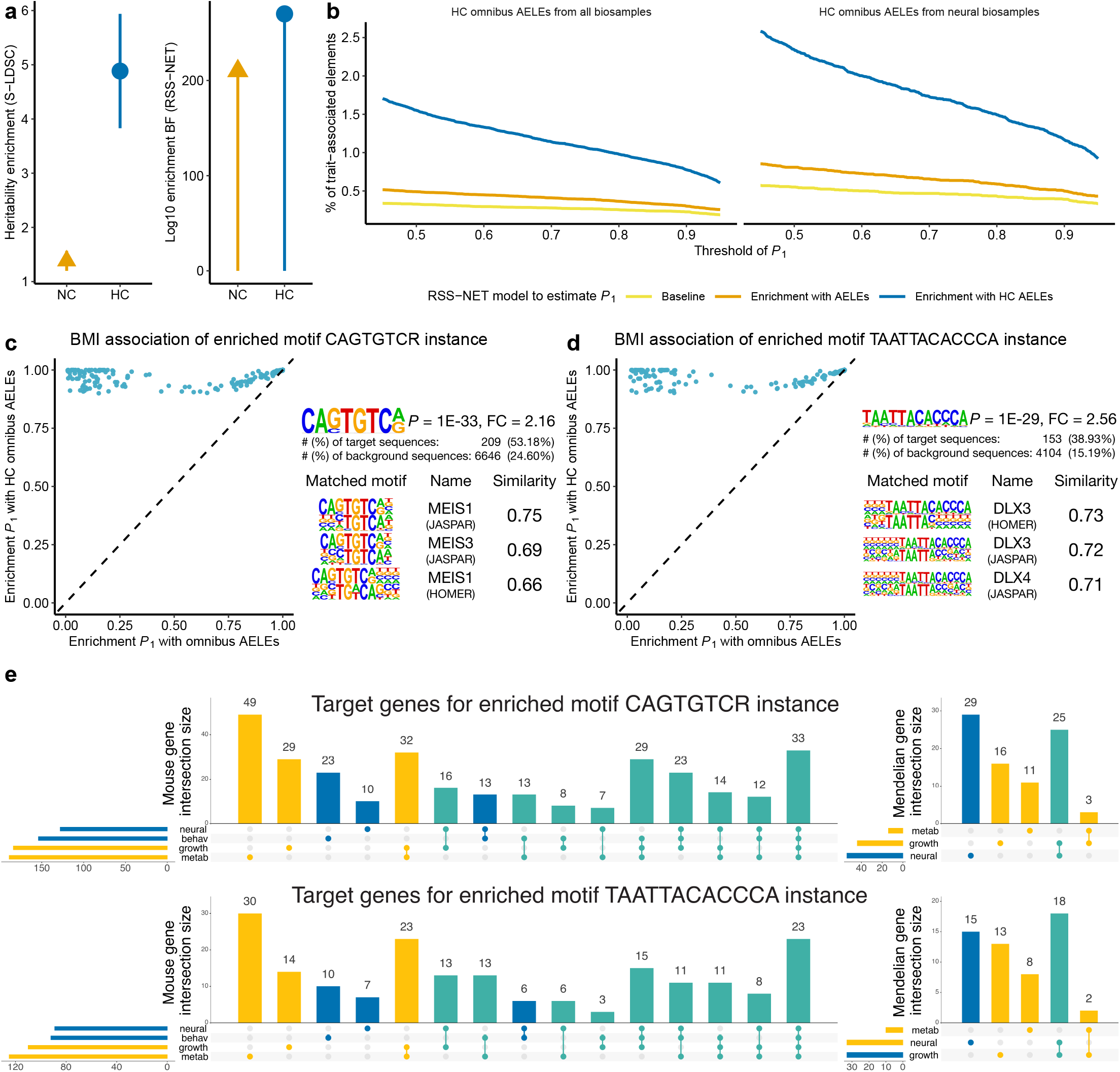
Prioritize HC AELEs for BMI. **a** Enrichments of heritability (S-LDSC) and genetic associations (RSS-NET) for omnibus AELEs without (NC) and with sequence conservation (HC). **b** Percentages of HC omnibus AELEs with 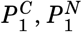, or 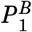 above a given threshold. BMI associations of **c** 210 HC AELEs that contain instances of the enriched motif CAGTGTCR and **d** 154 HC AELEs that contain instances of the enriched motif TAATTACACCCA. For **c**-**d**, each point denotes a HC ominbus AELE present in neural samples, with *y* and *x*-axis indicating its 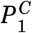 and 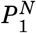 respectively. FC: fold change. **e** Overlap of the putative target genes (507 for CAGTGTCR-instance and 309 for TAATTACACCCA-instance HC AELEs respectively) with genes implicated in knockout mouse phenotypes and human Mendelian traits.

**Fig 6:**
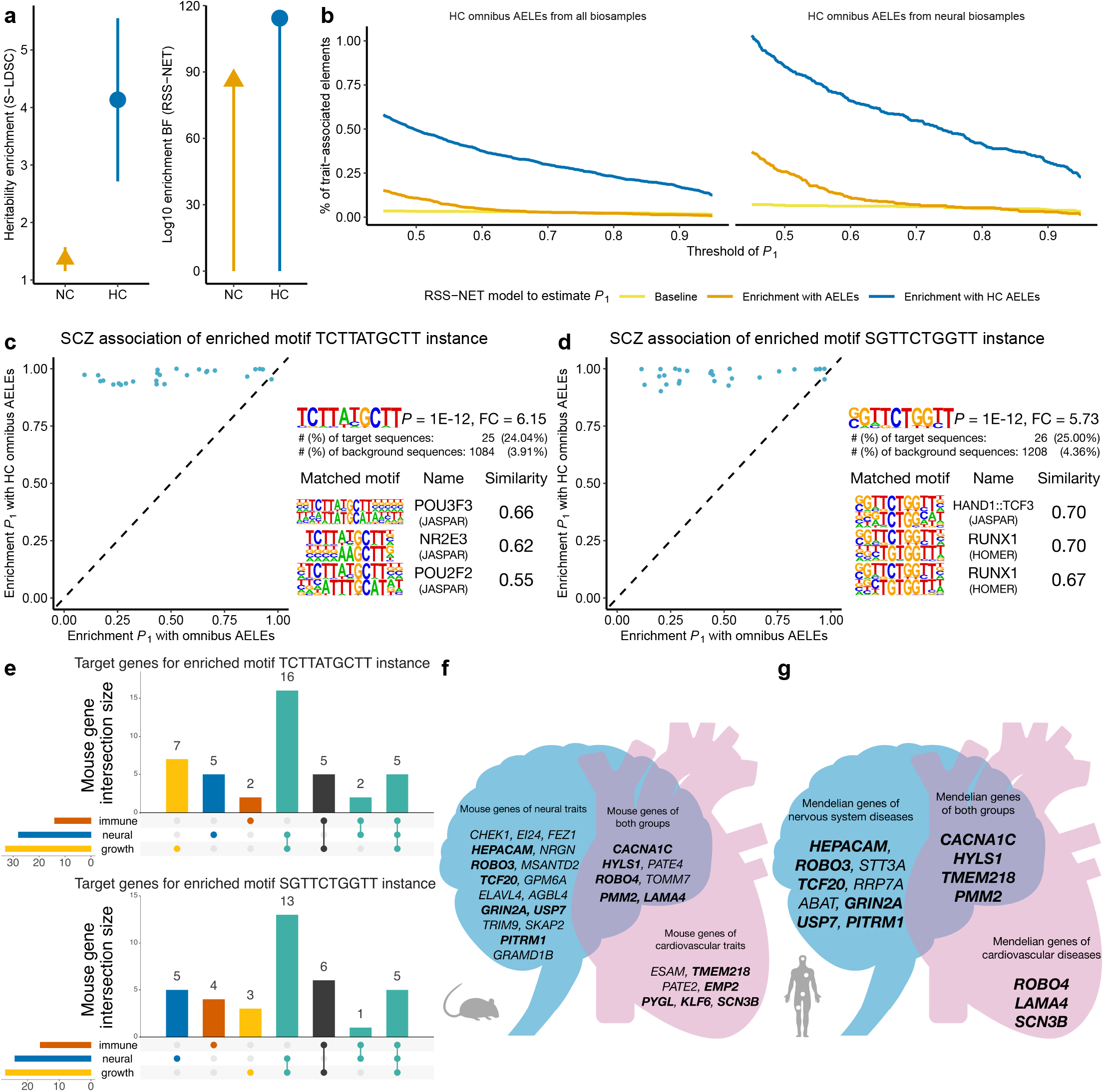
Prioritize HC AELEs for schizophrenia. Legends of **a**-**b** are the same as those in Fig. 5a-b. Schizophrenia associations of **c** 26 HC AELEs that contain instances of the enriched motif TCTTATGCTT and **d** 27 HC AELEs that contain instances of the enriched motif SGTTCTGGTT. For **c**-**d**, the rest is the same as Fig. 5c-d. **e** Overlap of the putative target genes (59 for TCTTATGCTT-instance and 59 for SGTTCTGGTT-instance HC AELEs) with genes implicated in knockout mouse phenotypes. Putative target genes of schizophrenia-associated HC AELEs with the enriched motif SGTTCTGGTT that cause **f** neural, cardiac or both types of knockout mouse phenotypes and **g** neural, cardiac or both types of human Mendelian traits.

Having confirmed their stronger enrichments, we focused on the genetic associations of HC omnibus AELEs. For each element, we computed its posterior probability that at least one SNP in this element is associated with the trait (*P*_1_), under the model where HC omnibus AELEs are enriched in associations with this trait (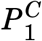; Methods). For comparison we also computed *P*_1_ for the same element-trait pair assuming (1) no enrichment 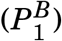 and (2) enrichment of omnibus AELEs 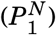. Consistent with previous work ^33,36^, the enrichment-informed 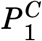 increases the inferred number of genetic associations (Table S15). Of 100591 HC omnibus AELEs, 781 are BMI-associated with 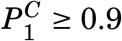, compared to 304 with 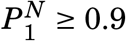 and 229 with 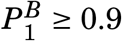 (Fig.5b). Similarly, 173 HC omnibus AELEs are associated with schizophrenia at 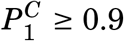, compared to 13 at 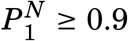 and 19 at 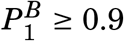 (Fig. 6b). When restricting to 33745 HC omnibus AELEs present in neural samples (a context highly relevant to BMI ^37^ and schizophrenia ^38^; Fig. 3b), we observed the same trend for BMI (394 with 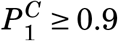, 168 with 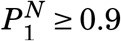, 132 with 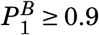) and schizophrenia (105 with 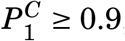, 10 with 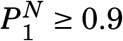, 14 with 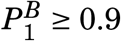. Together the results show that accounting for enrichment in HC omnibus AELEs enhances the strength of evidence for genetic associations (measured by *P*_1_), leading to potential discoveries missed by GWAS.

To assess regulatory functions underpinning identified associations, we searched for sequence motifs significantly enriched in a target set of traitassociated elements 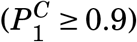 relative to a background set of non-associated elements 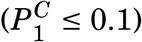. For both the target and background we only used HC omnibus AELEs present in neural samples (Fig. S1e-h) to minimize confounding introduced by sequence conservation or context specificity. We identified 127 and 3 enriched motifs from 394 BMI-associated (Table S16) and 105 schizophrenia-associated (Table S17) HC AELEs respectively. With the premise that HC AELEs affect a trait through downstream genes (Fig. S5), we further linked the trait-associated elements containing top-ranked motifs to putative target genes (Methods). Many linked genes are functionally relevant to the trait of interest (Fig. S6), as reported below.

### HC AELEs inform candidate effector genes for BMI

The 394 BMI-associated HC AELEs show the strongest enrichment of a sequence motif recognized by MEIS1 (Fig. 5c), which has key roles in adipogenesis ^39^ and neural development ^40^. Of 210 BMI-associated HC AELEs containing the MEIS1 motif, 107 are connected to 507 putative target genes (Table S18). Pathway analysis of these genes highlights multiple BMI-relevant processes, including apelin signaling, pituitary gland development and insulin secretion (Table S19). Though not implicated in GWAS ^34^, apelin and its receptors are involved in energy metabolism ^41^ and obesity ^42^.

The 394 BMI-associated HC AELEs are also strongly enriched for a motif of DLX3 (Fig. 5d), which is essential for vertebrate development ^43^. There are 154 BMI-associated HC AELEs containing the DLX3-binding motif, 74 of which are further linked to 309 putative target genes (Table S20). These genes are enriched in multiple processes related to body weight, such as pancreas development, apelin signaling and adipogenesis (Table S21).

To assess their biological and clinical themes, we looked up the 568 unique putative target genes in external databases (Table S22; Methods). Of the 568 genes, 407 and 146 are implicated in knockout mouse phenotypes and human Mendelian traits respectively. More than half of the 407 knockout mouse genes show growth (213), metabolic (217) and neural (228) phenotypes. A considerable fraction of the 146 Mendelian genes cause diseases characterized by growth (50), metabolic (17) and neural (59) phenotypes. Many of the neural genes are also related to growth and metabolism. Of 228 genes with neural mouse phenotypes, 181 (79.4%) have growth or metabolic phenotypes. Of 59 Mendelian genes with neural manifestations, 27 (45.8%) affect growth. We identified similar patterns when analyzing putative target genes informed by MEIS1 and DLX3-binding motifs separately (Fig. 5e). The gene results, together with neural enrichments of heritability (Fig. 3b) and fine-mapped variants (Fig. 4c), reinforce the key role of brain on body weight regulation ^37^.

Integrating BMI-associated HC AELEs with genes affecting mouse body weight and human monogenic obesity help prioritize effector genes for BMI. Apart from well-known obesity genes ^37^ (*LEP, PCSK1, NTRK2*), we identified several BMI effector genes that are supported by multiple converging lines of evidence but have not been reported in GWAS ^34^. For example, *CDK5* is a strong candidate for BMI: *CDK5* encodes cyclin-dependent kinase 5 (Cdk5) that has diverse functions ^44^ in neurons, adipocytes and beta cells; Cdk5 affects obesity and diabetes through phosphorylation of PPAR*γ* ^45^; and a mutation in *CDK5* causes lissencephaly with multiple neurodevelopmental features ^46^. *HSD11B1* is another plausible BMI effector: *HSD11B1* encodes 11*β*-hydroxysteroid dehydrogenase type 1 (11*β*-HSD1) that has key roles in obesity and related metabolic diseases ^47^; 11*β*-HSD1 overexpressed in adipose leads to visceral obesity and hyperphagia in mice ^48^; and mutations in *HSD11B1* affect the regeneration of cortisol ^49^, a steroid hormone associated with obesity ^50^. Further, *CDK5* and *HSD11B1* are therapeutic targets of a preclinical (L-751250) and Phase 1 (AZD8329) drugs for obesity respectively, and our results provide genetic support for the early-stage development.

### HC AELEs inform candidate effector genes for schizophrenia

The 105 schizophrenia-associated HC AELEs show strong enrichments of motifs recognized by POU3F3 (Fig. 6c) and HAND1::TCF3 complex (Fig. 6d), all of which are relevant to schizophrenia. POU3F3 is essential for cerebral cortex development ^51^. TCF3 regulates neocortical development through Wnt *β*-catenin signaling ^52^. HAND1 is critical for placenta development ^53^, which has been associated with the genetic risk of schizophrenia ^54^.

We identified 26 and 27 schizophrenia-associated HC AELEs containing POU3F3 and HAND1::TCF3-binding motifs respectively, further suggesting 86 unique putative target genes (Tables S23-S25). Of 31 genes linked to Mendelian diseases, 19 (61.3%) have neural indications. Among 59 genes with knockout mouse phenotypes available, many have immune (22), growth (43) and neural (38) phenotypes. Of 38 neural genes, 11 (28.9%) have immune and 29 (76.3%) have growth phenotypes in knockout mice. We observed similar neural-immune and neural-growth overlaps when analyzing POU3F3 and HAND1::TCF3 target genes separately (Fig. 6e). These findings recapitulate roles of immunity ^55^ and early development ^38^ in the etiology of schizophrenia.

Many putative target genes of schizophrenia-associated HC AELEs with the HAND1::TCF3-binding motif cause both neural and cardiac knockout mouse phenotypes (Fig. 6f) and human Mendelian traits (Fig. 6g), likely due to the key role of HAND1 in heart development ^53^. Our results highlight three genes (*CACNA1C, HYLS1, PMM2*) with neural-cardiac roles. *CACNA1C* has been repeatedly identified in GWAS of schizophrenia ^35^ and it causes arrhythmia associated with autism ^56^. Both *HYLS1* and *PMM2* have not been implicated in GWAS (Table S26), but their neural-cardiac roles are relevant to schizophrenia. *HYLS1* encodes hydrolethalus syndrome protein 1, which regulates the biogenesis and signaling of cilia ^57^. Cilia are antenna-like organelles with essential roles in cerebral cortical ^58^ and cardiac ^59^ development. A mutation in *HYLS1* causes hydrolethalus syndrome ^60^ characterized by developmental defects of fetal brain and heart. Mutations in *PMM2* cause a congenital disorder of glycosylation ^61^ with neurological and cardiac manifestations ^62^. Glycosylation has been linked to cardiovascular ^63^ and neuroinflammatory ^64^ diseases, as well as schizophrenia ^65^. In sum, HAND1::TCF3 target genes with neural-cardiac roles provide a means to elucidate genetic causes of comorbidity between schizophrenia and cardiovascular diseases ^66^.

## Discussion

Here we present a simple and scalable strategy to identify HC AELEs, active human enhancer-like elements that are highly conserved in the mouse genome, for 106 tissues and cell types. Combining them with 468 GWAS of EUR and EAS ancestries, we demonstrate that HC AELEs harbor a significant excess of genetic signals for human complex traits, as measured by common SNP heritability and fine-mapped variants. HC AELEs also help pinpoint previously undescribed but functionally relevant genes for BMI (e.g., *CDK5, HSD11B1*) and schizophrenia (e.g., *HYLS1, PMM2*). By integrating evolutionary and biochemical evidence, HC AELEs provide a unique perspective on genetic architecture, independent of existing functional annotations.

Our findings have important implications for complex trait genetics. First, compared to AELEs, HC AELEs have fewer sequence changes across species and stronger enrichments of trait heritability and causal variants, reinforcing the model of negative selection on genetic variants to affect complex traits ^13,23,35^. Second, HC AELEs capture consistent signals between EUR and EAS, highlighting the potential for cross-species methods like ours to improve the transferability across human populations ^29^. Third, though imperfectly conserved between humans and mice, HC AELEs retain regulatory functions to affect complex traits in a tissue-specific manner, corroborating the functional robustness of ultraconserved enhancers to mutations ^67^. Forth, HC AELEs highlight human sequences with high similarity in the mouse genome, suggesting a path to test human GWAS discoveries in mice.

Compared with previous efforts to integrate evolutionary and biochemical data, our study has several key strengths. First and foremost, exploiting a simple but profound idea of human-mouse sequence comparison to locate functional elements ^25,26^, our method can be easily scaled to hundreds of tissues as shown here, whereas existing studies often examine a single tissue ^18–23^. Second, unlike many studies that use either H3K27ac ^18,22,23^ or chromatin accessibility ^19–21^ alone to mark functional regions, we integrate both profiles to refine the annotation. Third, despite the human-mouse conservation, our method focuses firmly on functional sequences in the human genome. This approach contrasts with many studies ^18–21^ that focus on human orthologues of functional sequences in the mouse genome, bypassing the issue that the orthologues may not be functional in humans ^1^. Last but not the least, we assess conservation of genome sequences rather than H3K27ac signals across species ^22,23^. This choice not only yields significantly stronger heritability enrichments, but also eliminates the need for cross-species H3K27ac profiling in the same tissue, making the method more widely applicable.

Although tested on the mouse genome only, our pairwise comparison to identify HC AELEs is straightforward to implement more broadly for other species. Such extensions are feasible as high-quality genomes are becoming available for many species ^7,8^. That said, we caution that the pairwise approach might fall short in evolutionarily related species, such as humans and primates, due to the paucity of cross-species sequence variation. In such case, phylogentic modeling of multiple species may be worth pursuing ^3,16^. Another limitation is that sequence comparisons omit enhancers that are functionally conserved but nonorthologous at the sequence level ^68^. In such case, integrative modeling across functions and species may help ^21,69^. Despite the limitations, our method provides a useful benchmark for sophisticated models.

Currently HC AELEs are based on the bulk sequencing of H3K27ac and chromatin accessibility profiles, likely missing detailed cellular processes in which regulatory variants affect complex traits. Identifying cellularly resolved HC AELEs may be feasible, as single-cell epigenomic data are emerging. Indeed, single-cell atlases of chromatin accessibility have been recently established for many human tissues ^70^, and single-cell H3K27ac measurements will likely be available for diverse tissues soon with the advent of new technologies ^71^. Besides the single-cell extension, our current strategy may need to be adapted to incorporate other data such as chromatin conformation ^72,73^ and CRISPR screening ^32,74^ to capture the multifaceted nature of enhancers ^6^. Altogether, fine-tuning HC AELEs alongside advances in technologies and resources will markedly increase the resolution and accuracy.

Our findings emphasize the importance of combining evolutionary and biochemical evidence to understand the regulatory basis of heritable human traits. This integrative idea has been well documented and increasingly appreciated, but it remains underexploited at scale. This work represents a comprehensive effort and a major step forward to close this gap.

## Methods

### Reference genomes

We used the GRCh37 (hg19, human) and GRCm38 (mm10, mouse) genome assemblies throughout this study. We converted data based on GRCh38 (hg38) to GRCh37 using liftOver ^75^ (https://genome.ucsc.edu/cgi-bin/hgLiftOver) with the default setting and the minimum ratio of bases that must remap (‘minMatch’) being 0.95.

### Human epigenomes

We collected genome-wide sequencing data of 142 H3K27ac (ChIP-seq) and 171 chromatin accessibility (DNase-seq, ATAC-seq) profiles across 106 contexts (Table S1). We followed the ENCODE pipelines (https://www.encodeproject.org/data-standards) to identify H3K27ac and accessible chromatin peaks in each context.

### Sequence-conserved AELEs

To identify AELEs without sequence conservation (NC) for each of the 106 contexts, we first intersected its H3K27ac and accessible chromatin peaks using BEDTools ^76^ (version 2.27.1, https://bedtools.readthedocs.io). To identify LC, MC and HC AELEs for each context, we compared positions of NC AELEs between the human and mouse genomes using liftOver with the default setting and ‘minMatch’ as 0.1, 0.5 and 0.9, respectively (Fig. 1a). Each AELE defines an interval on the human genome, indicated by the chromosome, start and end positions. The codes and results are available at https://github.com/suwonglab/m2h-aele. Because the element counts of AELEs derived from an induced pluripotent stem (iPS DF 6.9) cell line are significantly lower than those in other contexts (NC: 547; LC: 387; MC: 360; HC: 182; Fig. 1b), we only used this cell line to create the omnibus version of AELEs and excluded it from all other analyses. To benchmark our primary approach (Fig. 1a), we developed an alternative method (Fig. S4a) to identify H3K27ac-conserved AELEs, leveraging 14 contexts that each had profiles of human chromatin accessibility, human and mouse H3K27ac available (Table S10). For each context, we first mapped H3K27ac peaks from the mouse to human genome using liftOver with the default setting and ‘minMatch’ being 0.9, and then we intersected the coordinate-converted mouse H3K27ac peaks with human H3K27ac and accessible chromatin peaks in the same context to create H3K27ac-conserved AELEs.

### SNP annotation

We stored each set of AELEs (omnibus or context-specific; without or with sequence conservation) as a BED file, consisting of one line per genomic interval (henceforth ‘element’). For each set of elements, we created the corresponding binary SNP annotation as:

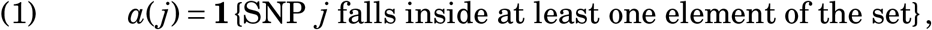

where *j* belongs to 5961159 EUR or 5469053 EAS common SNPs. Because omnibus AELEs aggregate all context-specific AELEs (Fig. S1a), we can show:

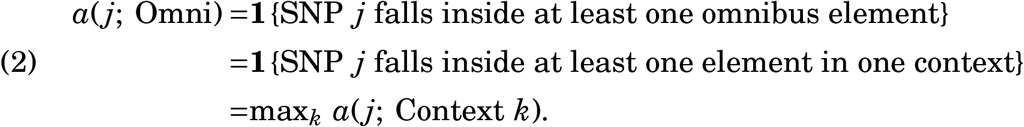

Equation (2) provides an alternative way to create the SNP annotation for omnibus AELEs directly from SNP annotations of context-specific AELEs, without identifying omnibus AELEs first and then applying Equation (1).

To assess the similarity of two binary SNP annotations (Fig. 1h-i), we computed Cramér’s *V* using the function ‘cramerV’ from R package rcompanion (version 2.4.16, https://cran.r-project.org/web/packages/rcompanion). To assess the similarity between a binary and a quantitative annotation (Fig. 1i), we computed Pearson’s *R* using the R built-in function ‘cor.test’.

### GWAS

We collected GWAS summary statistics from 312 UK Biobank GWAS, 108 EUR GWAS and 48 EAS GWAS (Table S5). The sample size of 468 datasets ranged from 14267 to 1320016, with a median of 452264. All datasets had observed-scale heritability *Z*-scores ≥6 as estimated by S-LDSC (see below). All datasets were processed as previously described ^13,23^.

### Heritability enrichment

To assess the heritability enrichment of a SNP annotation in a GWAS, we used S-LDSC ^13^ (version 1.0.1, https://github.com/bulik/ldsc) with 1000 Genomes ^77^ Phase 3 as the LD reference panel (9997231 EUR and 8768561 EAS reference SNPs) and 96 annotations from the baselineLD model ^23^ as covariates (version 2.2, https://alkesgroup.broadinstitute.org/LDSCORE). The 96 annotations capture diverse functions in the genome such as translation, regulation and selection (Table S4). To analyze (omnibus or context-specific) AELEs without sequence conservation in a GWAS, we modeled the variance of effect size for SNP *j* as

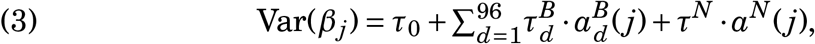

where *τ*_0_ is the background per-SNP contribution to heritability, 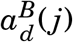 is the value of SNP *j* for one of the 96 baseline annotations, *a*^*N*^(*j*) = 1 if SNP *j* falls inside any AELE and 0 otherwise, and 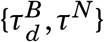 are per-SNP contributions of one unit of the corresponding annotations to heritability. Equation (3) allows us to assess the contribution of AELEs to heritability conditional on 96 known annotations, which helps reduce bias due to model misspecification ^13,23^.

To analyze an annotation of (omnibus or context-specific) AELEs with sequence conservation (LC, MC, HC) in a GWAS, we extended Equation (3) as

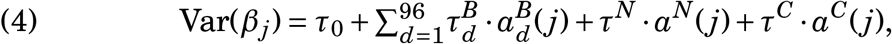

where *a*^*C*^(*j*) 1 if SNP *j* falls inside any conserved AELE and 0 otherwise, and *τ*^*C*^ is the per-SNP contribution of one unit of the conserved AELE annotation to heritability. Like Equation (3), this model captures the unique contribution of conserved AELEs to heritability conditional on 96 known annotations and AELEs without sequence conservation in the same context.

We used two quantities ^13,23^ to summarize S-LDSC results. First, we computed the heritability enrichment of an annotation *a* in a GWAS as

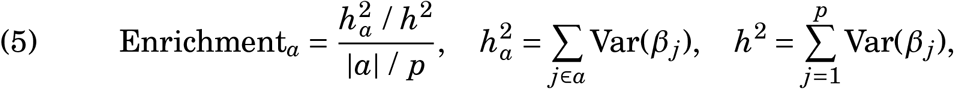

where |*a*| is the number of common SNPs with annotation *a, p* is the total number of common SNPs (*p* = 5961159 for EUR and 5469053 for EAS), and 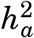 and *h*^2^ are heritabilities due to |*a*| common SNPs with annotation *a* and *p* common SNPs respectively. Second, we computed the standardized effect size (*τ*^⋆^) of an annotation *a* in a GWAS as

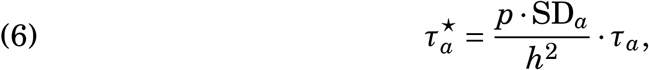

where SD_*a*_ is the standard deviation of annotation *a* across *p* common SNPs and *τ*_*a*_ is the original effect size for annotation *a* in Equations (3)-(4). Both quantities can be compared across GWAS and annotations. Unlike the enrichment in Equation (5), *τ*^⋆^ in Equation (6) can capture the unique effect of annotation *a* conditional on all other annotations in Equations (3)-(4).

To meta-analyze S-LDSC results across traits and contexts, we used the function ‘meta.summaries’ from R package rmeta (version 3.0, https://cran.r-project.org/web/packages/rmeta) as previously described ^13,23^. For both heritability enrichment and *τ*^⋆^, we performed random-effects meta-analyses of individual estimates and SEs to obtain meta-analyzed estimates and SEs (Fig.s 2a-b, 3a-b, S2-S4). To find the *P*-value for meta-analyzed heritability enrichment, we first meta-analyzed 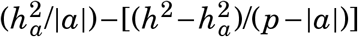 and then computed a one-sided *Z*-score to test if this difference is greater than 0. To find the *P*-value for meta-analyzed *τ*^⋆^, we computed a one-sided *Z*-score to test if the meta-analyzed estimate is greater than 0.

To assess concordance of S-LDSC results between EUR and EAS GWAS of the same trait, we use the function ‘deming’ from R package deming (version 1.4, https://cran.r-project.org/web/packages/deming) as previously described ^29^. For each annotation, we fitted a generalized Deming regression of EUR estimates on EAS estimates across 34 traits, while accounting for SEs.

### Fine mapping

The fine-mapping results ^32^ of 94 traits in UK Biobank were produced by FINEMAP ^78^ and SUSIE ^79^ (version 1.1, https://www.finucanelab.org/data). Here we excluded variants without any 95% credible set assigned, variants in LD (*R*^2^ > 0.6) with a variant failing Hardy Weinberg equilibrium test (*P* < 1 × 10^−12^), and variants in LD (*R*^2^ > 0.8) with a common EUR structural variant. We further intersected the variants with 9997231 EUR SNPs in 1000 Genomes, yielding a final set of 515848 unique SNPs for this study.

### Trait-associated HC AELEs

We previously developed RSS-NET ^33^ that integrates GWAS summary statistics with genomic annotations to identify genetic enrichments and associations simultaneously. Here we expanded this Bayesian framework to prioritize trait-associated HC AELEs. Specifically, we combined the RSS likelihood ^80^ with the following prior distribution:

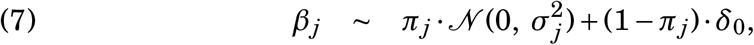

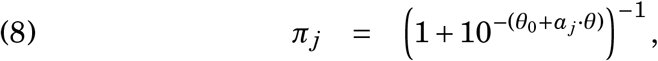

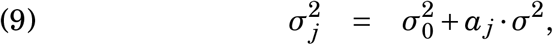

where *β*_*j*_ denotes the effect of SNP *j* on a given trait, *π*_*j*_ denotes the probability that SNP *j* is associated with the trait (*β*_*j*_ ≠ 0), 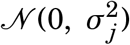 denotes a normal distribution with mean 0 and variance 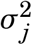 specifying the effect size of a trait-associated SNP *j, δ*_0_ denotes point mass at zero (*β*_*j*_ = 0), and *a*_*j*_ = 1 if SNP *j* falls inside any HC AELE and 0 otherwise. In Equation (8), the baseline parameter *θ*_0_ < 0 captures the genome-wide background fraction of trait-associated SNPs, and the enrichment parameter *θ* > 0 reflects the increase in probability that a SNP inside HC AELEs is trait-associated. In Equation (9), the baseline parameter 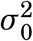 captures the genome-wide background effect size of trait-associated SNPs, and the enrichment parameter *σ*^2^ reflects the increase in effect size of trait-associated SNPs inside HC AELEs. We specified hyper-priors on the unknown parameters 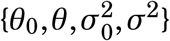 and used variational nference to compute posterior distributions as previously described^33,36^. The implementation is available at https://github.com/suwonglab/rss-net.

To assess whether HC AELEs are enriched for genetic associations with a target trait (Fig.s 5a, 6a), we computed a Bayes factor (BF):

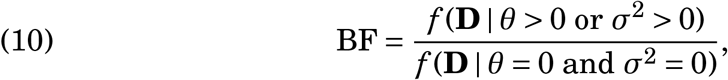

Where *f*(·) denotes the marginal likelihood for the extended RSS-NET model and **D** is a shorthand for the input data including GWAS summary statistics, LD estimates and SNP annotations of HC AELEs (*a*_*j*_). A larger BF indicates stronger evidence for enrichment of genetic associations.

To identify if a HC AELE is associated with a trait, we used *P*_1_, the posterior probability that at least one SNP in the HC AELE is trait-associated:

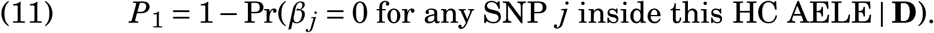

For each HC AELE, 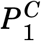 (*y*-axis of Fig.s 5c-d, 6c-d), 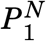 (*x*-axis of Fig.s 5c-d, 6c-d) and 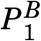 are *P*_1_ values evaluated with different definitions of *a*_*j*_. For 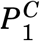, *a*_*j*_ = 1 if SNP *j* falls inside any HC AELE and 0 otherwise. For 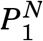, *a*_*j*_ = 1 if SNP *j* falls inside any AELE and 0 otherwise. For 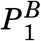, *a*_*j*_ = 0 for any SNP *j* (i.e., no enrichment), which is equivalent to *θ* = *σ*^2^ = 0 in Equations (8)-(9).

We applied the RSS-NET extension to EUR GWAS of BMI ^34^ (https://portals.broadinstitute.org/collaboration/giant) and schizophrenia ^35^ (https://walters.psycm.cf.ac.uk) as previously described ^33^. For each GWAS, we analyzed both omnibus and HC omnibus AELEs. We did not analyze HC context-specific AELEs with RSS-NET because they contain less than 0.45% of common SNPs (Fig. 1g), leading to sparse SNP annotations (i.e., *a*_*j*_ = 0 for most SNPs). Reliable estimation of the RSS-NET enrichment parameters (*θ, σ*^2^) in Equations (8)-(9) requires sufficient SNPs with *a*_*j*_ = 1.

### Motif enrichment

We used the HOMER ^81^ command ‘findMotifsGenome.pl’ (version 4.11, http://homer.ucsd.edu/homer/) to identify genomic regions specifically enriched in a target set of sequences against a background set. We used the exact regions provided (‘-size given’) and searched for known motifs (‘-mknown’) in the curated list ^72,82^ of 1465 unique motifs (https://github.com/suwonglab/peca). Beyond the 1465 known motifs, we also identified de novo motifs, and matched them to known motifs based on similarity of motif matrices (Fig.s 5c-d, 6c-d). For each motif we computed a fold change of fractions of target against background sequences containing the motif, and a binomial *P*-value to quantify enrichment. We identified a significant enrichment of a motif when the fold change ≥ 2 and *P* ≤ 1 × 10^−12^.

### Target genes of HC AELEs

To link trait-associated HC AELEs to putative target genes, we used cCRE-gene linkages derived from adult mouse cerebrum ^20^, human fetal ^83^ and adult ^84^ brain. We used BEDTools to overlap HC AELEs with cCREs in these datasets and then used the cCRE-gene linkages to identify putative target genes for HC AELEs.

The adult mouse dataset (http://catlas.org/mousebrain) contains 813638 linkages that connect 261204 cCREs to 12722 putative target genes. We first converted HC AELEs (GRCh37) to regions in the mouse genome (GRCm38), overlapped them with the mouse cCREs to find the linked mouse genes, and then converted the MGI ^85^ gene symbols to HGNC symbols.

The human fetal dataset (https://doi.org/10.6084/m9.figshare.16829101.v1) provides 63653 linkages of 33862 cCREs and 10892 genes for cortical plate, and 63740 linkages of 34044 cCREs and 11146 genes for germinal zone. The human adult dataset (http://resource.psychencode.org) is based on the repeat-masked transcription factor binding site map (GRN1) and the complete map (GRN2). GRN1 has 577529 linkages of 71097 cCREs and 13308 genes. GRN2 has 531322 linkages of 70532 cCREs and 13330 genes.

### External databases

To interpret our findings, we used the following external databases: phastCons ^16^ 100-vertebrate scores (http://hgdownload.cse.ucsc.edu/goldenPath/hg38/phastCons100way), ENCODE ^9^ human cCREs (version 3, https://screen.encodeproject.org), Metascape ^86^ for biological pathways (version 3.5, https://metascape.org), MGI ^85^ for knockout mouse genes (version 6.18, http://www.informatics.jax.org), OMIM ^87^ for human Mendelian genes (https://www.omim.org), TTD ^88^ for drug target genes (version 8.1.01, https://idrblab.org/ttd) and GWAS Catalog ^89^ for GWAS-implicated genes (version 1.0.2, https://www.ebi.ac.uk/gwas).

## Data and code availability

The AELEs (omnibus or context-specific, without or with sequence conservation) and source codes are available at https://github.com/suwonglab/m2h-aele. Links and identifiers of other data and codes are specified in Tables S1, S5, S10 and Methods.

## Acknowledgments

X.Z. is supported by Stein Fellowship from Stanford University, Institute for Computational and Data Sciences Seed Grant and Consortium on Substance Use and Addiction Seed Grant from Pennsylvania State University. W.H.W is supported by NIH grants P50HG007735 and R01HG010359 and NSF grant DMS1952386. This study uses resources at the Stanford Research Computing Center.

## Author contributions

X.Z. and W.H.W. conceived and supervised the study. S.M. collected the epigenomic data. X.Z. collected the GWAS data. S.M. generated the context-specific AELEs and linkages of target genes. X.Z. generated the omnibus AELEs and results related to GWAS. X.Z. performed the statistical analyses and database lookups. X.Z. created the figures and supplemental materials. X.Z. wrote the manuscript. All authors reviewed the manuscript.

## Supplementary table legends

**Table S1: Human epigenomic data**. Accession numbers of H3K27ac and chromatin accessibility profiles for 106 human tissues and cell types.

**Table S2: Motif enrichments of AELEs**. For each set of AELEs (omnibus or context-specific, without or with sequence conservation), significant motif enrichments are identified by HOMER (≥ 2-fold and *P* ≤ 1 × 10^−12^). For AELEs without conservation (NC), the background is set as randomly selected sequences that match the GC-content distribution of NC AELEs. For AELEs with conservation (LC, MC, HC), the background is set as either (1) GC-matched random sequences or (2) NC AELEs in the same context.

**Table S3: Pairwise correlations for AELE annotations of SNPs**. For each set of AELEs (omnibus or context-specific, without or with sequence conservation), the binary SNP annotation is created for both EUR and EAS populations. For each population, two sets of genome-wide SNPs are used: ‘all SNPs’ (EUR: 9997231; EAS: 8768561); ‘common SNPs’ (EUR: 5961159; EAS: 5469053). For each pair of SNP annotations, the correlation is measured by Cramér’s *V*, which ranges from 0 (no association) to 1 (perfect association).

**Table S4: Correlations between AELE annotations and 96 existing annotations of SNPs**. The S-LDSC baselineLD model consists of 83 binary and 13 quantitative annotations of SNPs. The correlation between an AELE annotation and a binary S-LDSC annotation is measured by Cramér’s *V*. The correlation between an AELE annotation and a quantitative S-LDSC annotation is measured by Pearson’s *R*. The rest is the same as Table S3.

**Table S5: GWAS data**. Phenotype descriptions, publications (PubMed IDs) and sample sizes for 468 GWAS datasets.

**Table S6: Heritability enrichments of omnibus AELEs**. S-LDSC results of SNP annotations based on omnibus AELEs without and with sequence conservation (LC, MC, HC) for 468 GWAS datasets.

**Table S7: Meta-analyzed heritability enrichments of omnibus AELEs**. S-LDSC results of SNP annotations based on omnibus AELEs without and with sequence conservation (LC, MC, HC) in 30 strata.

**Table S8: Heritability enrichments of HC context-specific AELEs**. SLDSC results of HC context-specific AELEs for 468 GWAS and 105 contexts.

**Table S9: Meta-analyzed heritability enrichments of HC context-specific AELEs**. S-LDSC results of HC context-specific AELEs in 432 strata.

**Table S10: Mouse epigenomic data**. Accession numbers of H3K27ac profiles for 14 mouse contexts, and context-matching human epigenomic data.

**Table S11: Heritability enrichments of AELEs with H3K27ac conservation**. S-LDSC results of SNP annotations based on HC context-specific AELEs using human-mouse H3K27ac comparison and genome comparison respectively, and NC context-specific AELEs, for 379 GWAS and 14 contexts.

**Table S12: Meta-analyzed heritability enrichments of AELEs with H3K27ac conservation**. S-LDSC results of HC AELEs using human-mouse H3K27ac comparison and genome comparison respectively, for 34 strata.

**Table S13: Fractions of fine-mapped SNPs in omnibus AELEs**. Each row reports the following fraction:

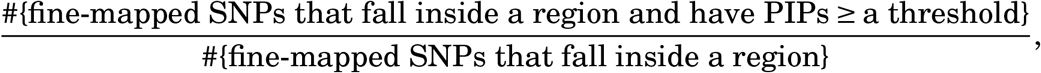

for a given combination of GWAS dataset, fine-mapping method and PIP threshold (columns 1-3), where the ‘region’ denotes the whole genome, omnibus AELEs without or with sequence conservation (columns 4-8).

**Table S14: Fractions of fine-mapped SNPs in HC context-specific AELEs**. Each row reports the following fraction (column 6):

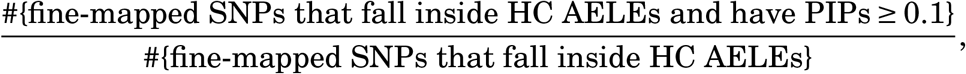

for a given combination of GWAS dataset, fine-mapping method and context group (columns 1-3), together with a binomial *P*-value (column 7) testing if this fraction is greater than the fraction of fine-mapped SNPs genome wide that have PIPs ≥ 0.1 (using the same GWAS data and fine-mapping method).

**Table S15: Genetic associations of HC omnibus AELEs with BMI and schizophrenia**. Each row represents a HC omnibus AELE (columns 1-3), and reports the estimated 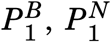 and 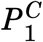 values based on GWAS of BMI (columns 5-7) and schizophrenia (columns 8-10). The 4th column (‘Neural’) indicates if the element contains any HC context-specific AELE derived from neural samples (1: yes; 0: no), as visualized in Fig. S1e-h.

**Table S16: Motif enrichments of AELEs associated with BMI**. Each row represents a de novo or known motif whose enrichment result is generated by HOMER. The target sequences are HC omnibus AELEs that are present in neural samples and have BMI-associated 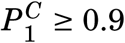, as indicated in columns 4 and 7 of Table S15. The background sequences are HC omnibus AELEs that are present in neural samples and have BMI-associated 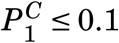.

**Table S17: Motif enrichments of AELEs associated with schizophrenia**. The target and background sequences are neural-related HC omnibus AELEs with schizophrenia-associated 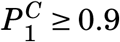 and 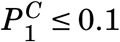 respectively (see columns 4 and 10 of Table S15). The rest is the same as Table S16.

**Table S18: Putative target genes of BMI-associated HC AELEs with MEIS1-binding motif**. Each row reports a published linkage between a neural-related, BMI-associated HC omnibus AELE 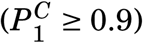 that contains the enriched sequence motif recognized by MEIS1 and a likely target gene, together with the corresponding publication (PubMed ID).

**Table S19: Pathway enrichments of target genes in Table S18**. Each row reports a Metascape pathway (columns 1-3), the number and fraction of target genes that belong to this pathway (columns 4-5), the log10 *P*-value (column 6) based on the hypergeometric distribution (with all genes in the genome as the enrichment background), and the log10 *q*-value (column 7) based on the Benjamini-Hochberg procedure to account for multiple testing.

**Table S20: Putative target genes of BMI-associated HC AELEs with DLX3-binding motif**. All elements listed here contain instances of the enriched DLX3-binding motif. The rest is the same as Table S18.

**Table S21: Pathway enrichments of target genes in Table S20**. The legend is the same as the legend of Table S19.

**Table S22: External database lookup of genes in Tables S18 and S20**. Each row reports identifiers of a gene in Mouse Genome Informatics (MGI), Online Mendelian Inheritance in Man (OMIM) and Therapeutic Target Database (TTD). A blank entry indicates the absence of a gene in a database.

**Table S23: Putative target genes of schizophrenia-associated HC AELEs with POU3F3-binding motif**. The elements listed here are neuralrelated, schizophrenia-associated HC omnibus AELEs 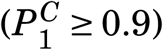 that contain the enriched motif recognized by POU3F3. The rest is the same as Table S18.

**Table S24: Putative target genes of schizophrenia-associated HC AELEs with HAND1::TCF3-binding motif**. The elements listed here are neuralrelated, schizophrenia-associated HC omnibus AELEs 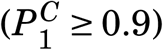 that contain the enriched HAND1::TCF3 motif. The rest is the same as Table S18.

**Table S25: External database lookup of genes in Tables S23 and S24**. The legend is the same as Table S22.

**Table S26: GWAS Catalog lookup**. The column heading description is available at https://www.ebi.ac.uk/gwas/docs/fileheaders.

**Fig S1:**
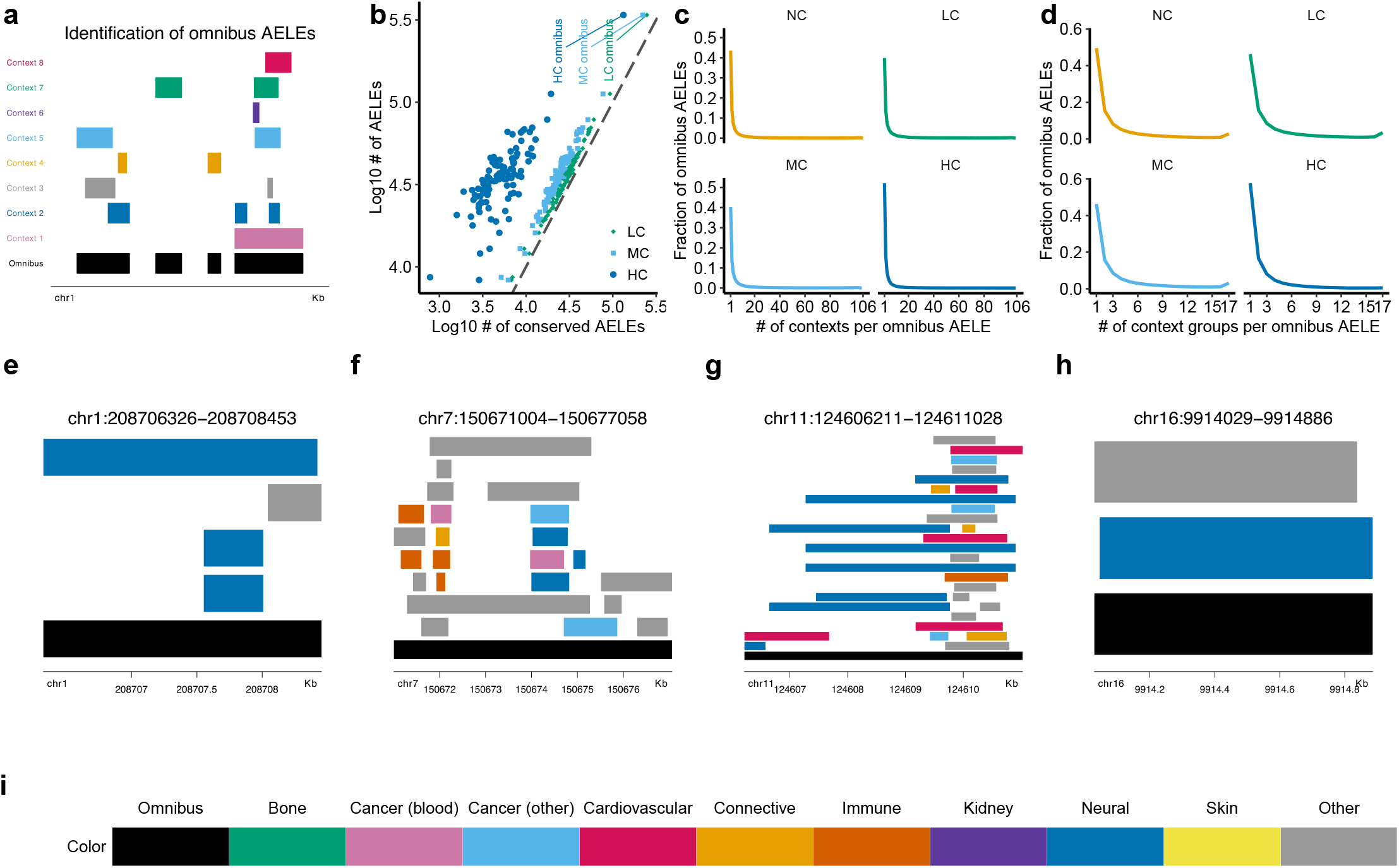
Identification and characterization of omnibus AELEs. **a** Context-specific AELEs from various contexts (colored bars) are first aggregated and then the overlapping segments are merged into 4 non-overlapping omnibus AELEs (black bars). **b** Element counts for context-specific and omnibus AELEs without (*y*-axis) and with (*x*-axis) sequence conservation. Distributions of numbers of **c** contexts and **d** context groups underlying each omnibus AELE, without (NC) and with (LC, MC, HC) sequence conservation. Illustration of 4 neural-related omnibus AELEs that contain significant genetic associations with **e**-**f** BMI (Fig. 5) and **g**-**h** schizophrenia (Fig. 6). **i** Color legends for context groups shown in **e**-**h**, as well as Fig.s 3 and S3.

**Fig S2:**
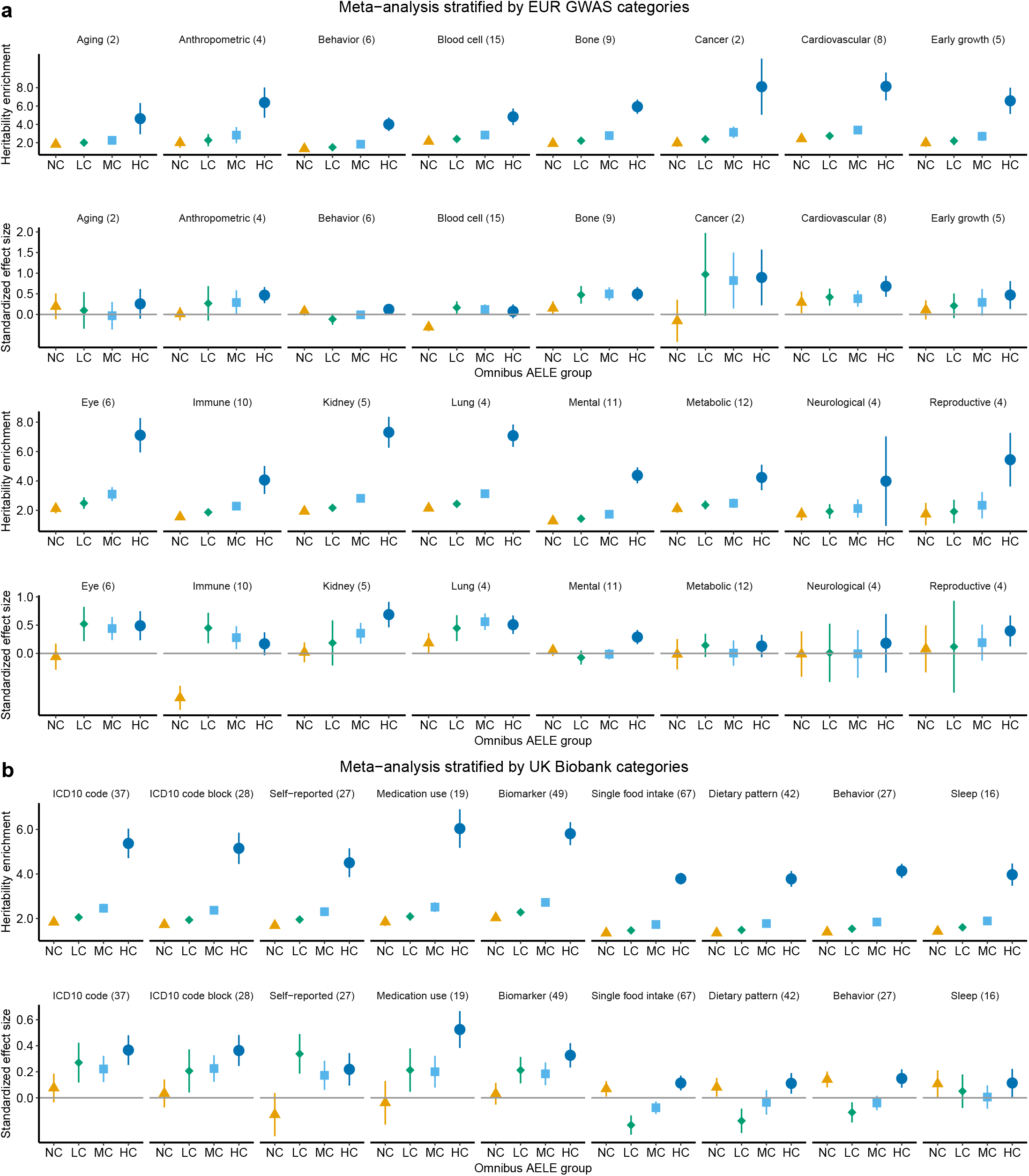
Additional results of heritability enrichments for omnibus AELEs with varying levels of sequence conservation. Estimates meta-analyzed within each of **a** 16 trait groups for the 108 EUR GWAS datasets and **b** 9 trait groups for the 312 UK Biobank datasets. Numerical results are available in Table S7. The legend is the same as Fig.s 2a-b.

**Fig S3:**
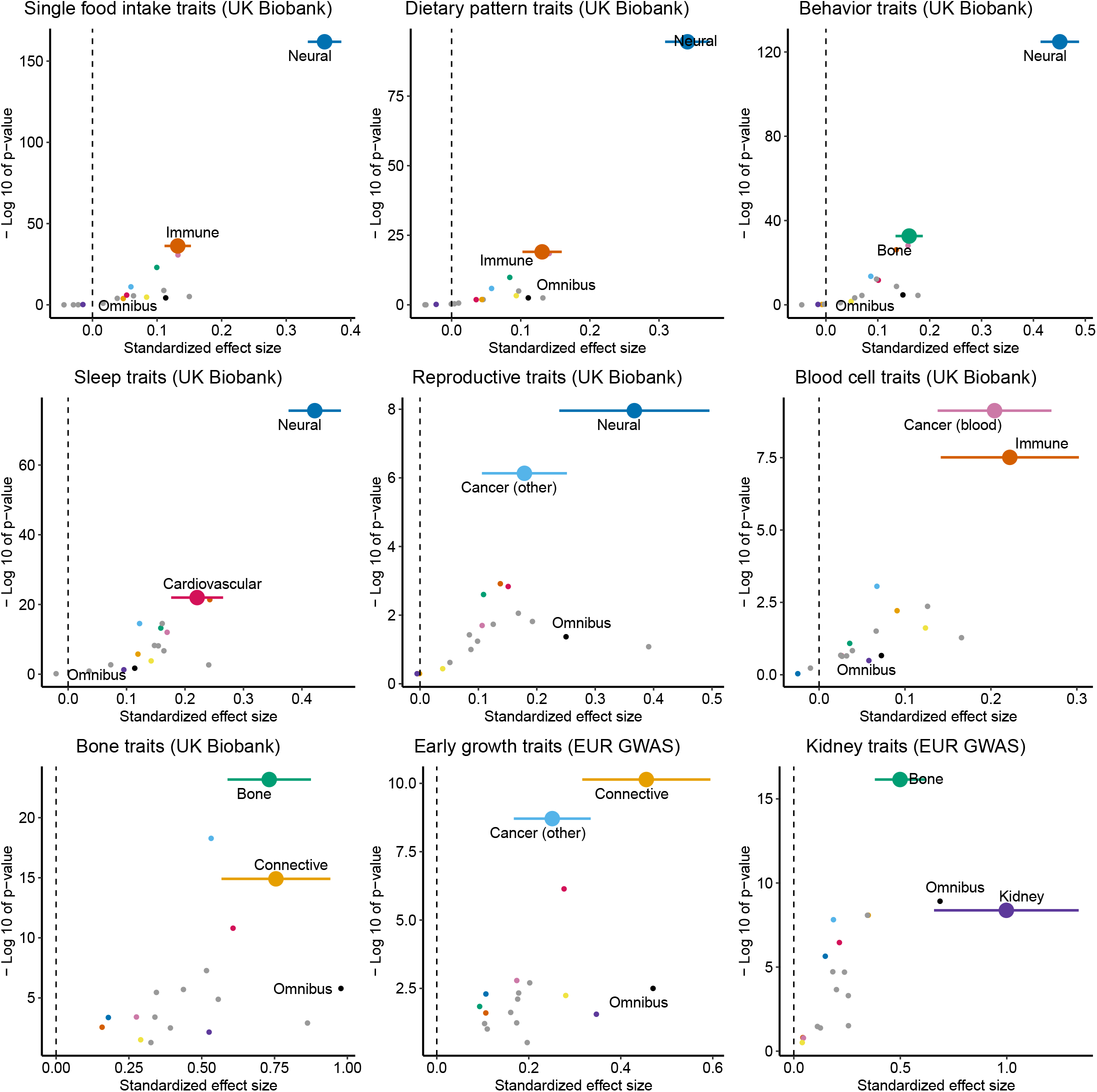
Additional results of heritability enrichments for HC context-specific AELEs. The legend is the same as Fig. 3a. For behavior, blood cell and bone traits, the results shown in Fig. 3a are based on the EUR GWAS datasets whereas the results shown here are based on the UK Biobank datasets.

**Fig S4:**
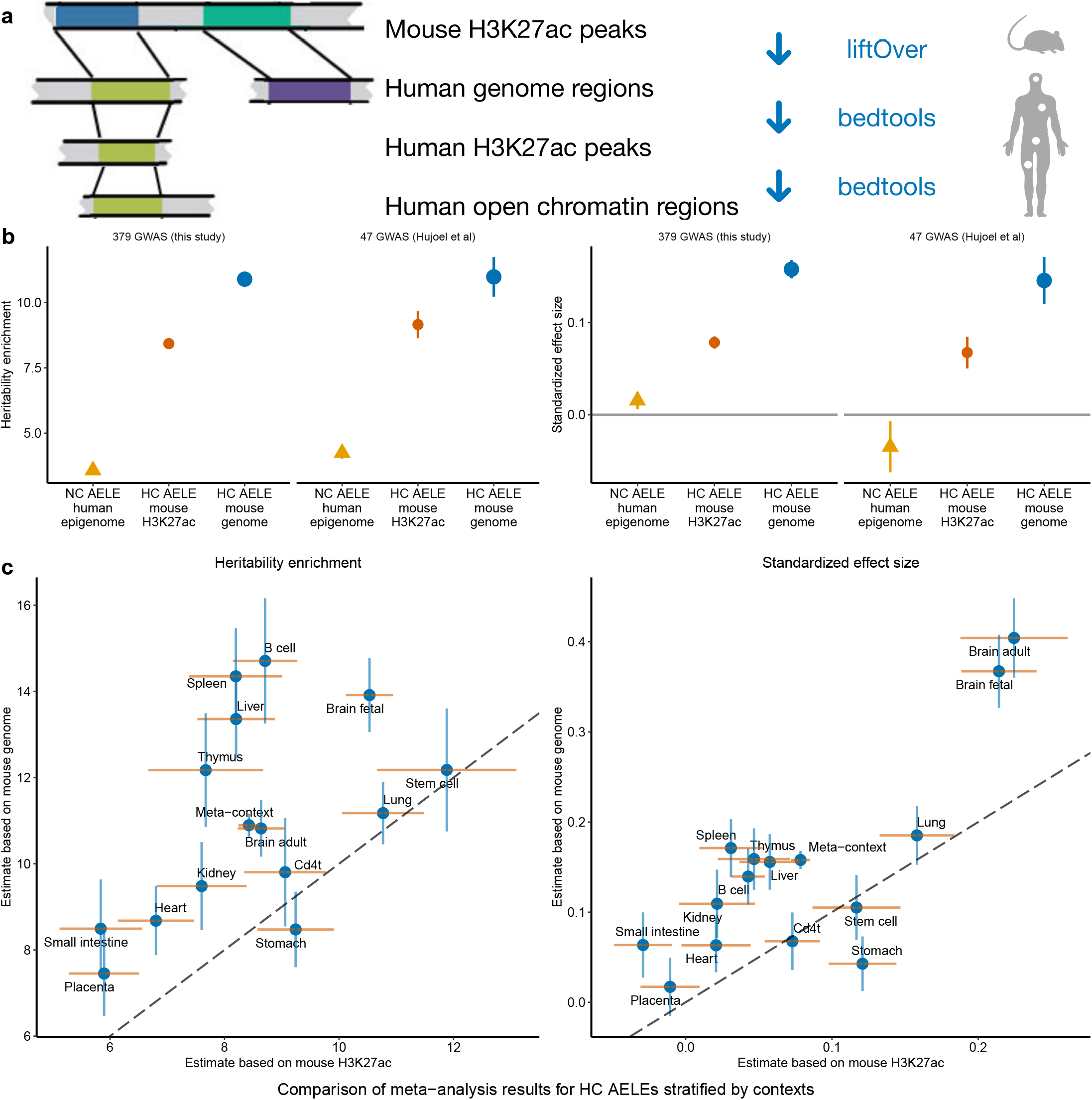
Comparison of common SNP heritability enrichments for HC AELEs based on the mouse genome and H3K27ac profile. For each trait and each of the 14 contexts with mouse H3K27ac ChIP-seq data available (Table S10), we estimate the heritability enrichment and standardized effect size (*τ*^⋆^) for HC AELEs based on the conservation with the mouse genome (Fig. 1a) and mouse H3K27ac profile, both conditioning on AELEs (NC) in the same context and 96 known annotations. **a** Schematic of creating HC AELEs based on the mouse H3K27ac profile. **b** Estimates meta-analyzed across 14 contexts and 379 GWAS datasets curated for this study, or a previously defined set of 47 independent GWAS datasets. **c** Estimates meta-analyzed across 379 GWAS datasets for each of the 14 contexts. The dashed lines have intercept 0 and slope 1. Numerical results of **b** and **c** are available in Table S12. The ‘meta-context’ results in **c** correspond to the meta-analyzed results of 379 GWAS in **b**. For **b**-**c**, each point denotes an estimate and each error bar denotes ±1.96 SE.

**Fig S5:**
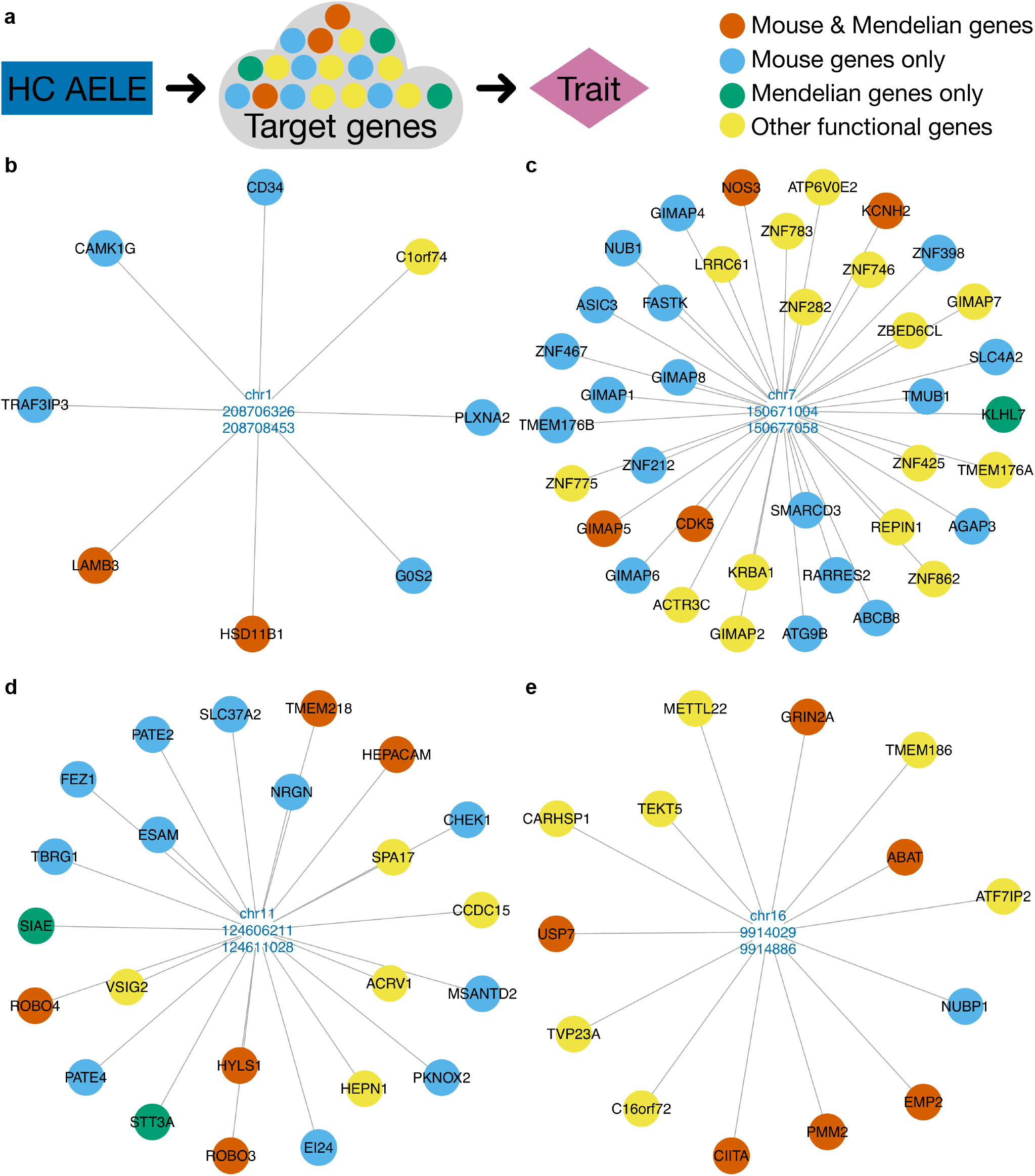
Putative target genes of BMI and schizophrenia-associated HC AELEs. **a** HC AELEs affect a complex trait through regulating target genes with functional impact on the trait. Putative target genes (Table S18) of two BMI-associated HC AELEs that contain instances of the enriched motif CAGTGTCR (Fig. 5c): **b** chr1:208706326-208708453 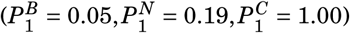, **c** chr7:150671004-150677058 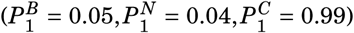. Putative target genes (Table S24) of two schizophrenia-associated HC AELEs that contain instances of the enriched motif SGTTCTGGTT (Fig. 6d): **d** chr11:124606211-124611028 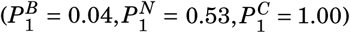, **e** chr16:9914029-9914886 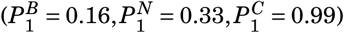.

**Fig S6:**
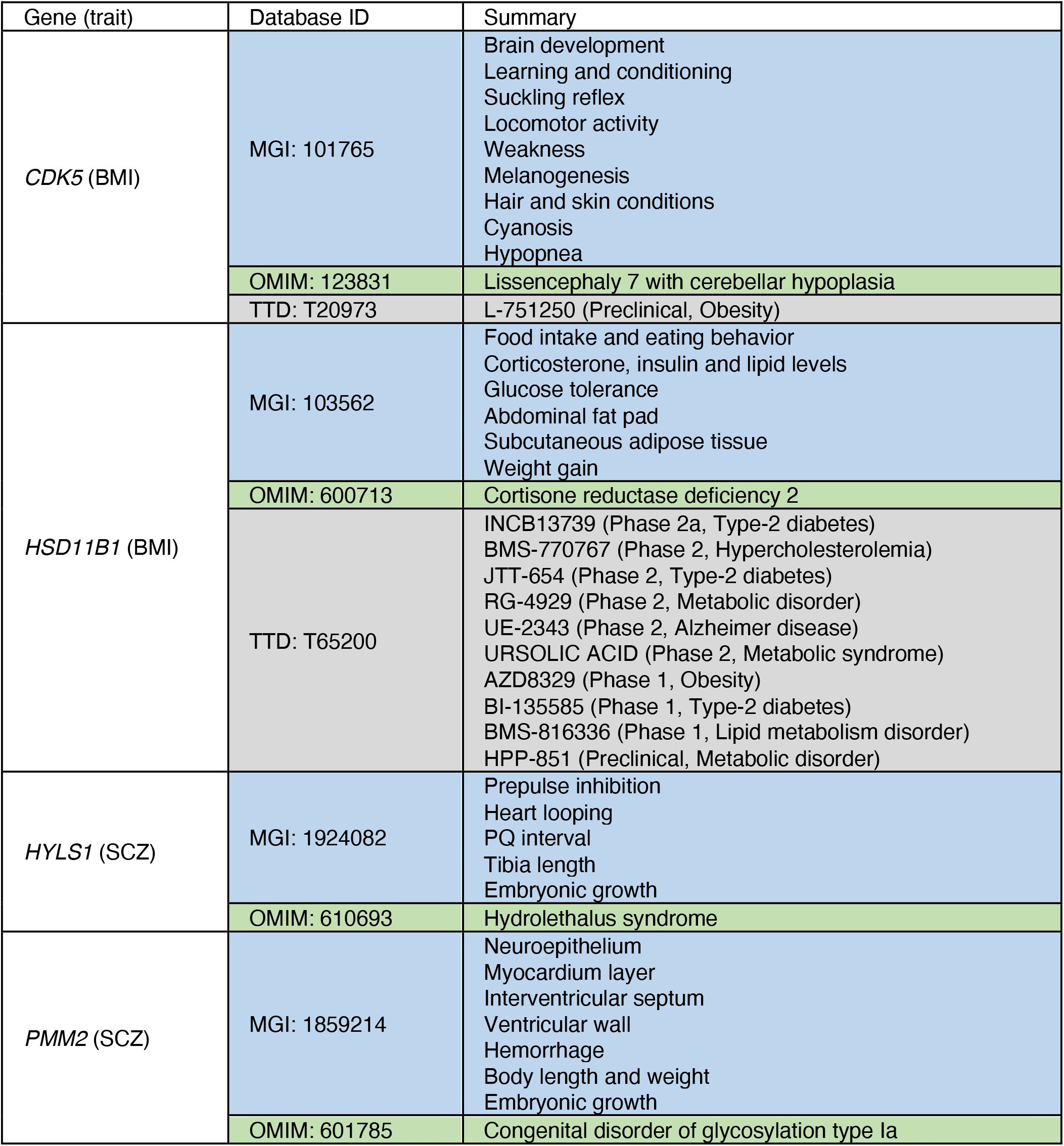
External database lookup of select downstream genes of BMI and schizophrenia-associated HC AELEs. BMI: body mass index. SCZ: schizophrenia. MGI: Mouse Genome Informatics. OMIM: Online Mendelian Inheritance in Man. TTD: Therapeutic Target Database. The lookups of all putative target genes are provided in Tables S22 and S25.

## References

[1] Kellis, M. et al. Defining functional DNA elements in the human genome. Proc. Natl. Acad. Sci. U.S.A. 111, 6131–6138 (2014).

[2] Loots, G. G. et al. Identification of a coordinate regulator of interleukins 4, 13, and 5 by cross-species sequence comparisons. Science 288, 136–140 (2000).

[3] Boffelli, D. et al. Phylogenetic shadowing of primate sequences to find functional regions of the human genome. Science 299, 1391–1394 (2003).

[4] Visel, A. et al. ChIP-seq accurately predicts tissue-specific activity of enhancers. Nature 457, 854–858 (2009).

[5] Creyghton, M. P. et al. Histone H3K27ac separates active from poised enhancers and predicts developmental state. Proc. Natl. Acad. Sci. U.S.A. 107, 21931–21936 (2010).

[6] Gasperini, M., Tome, J. M. & Shendure, J. Towards a comprehensive catalogue of validated and target-linked human enhancers. Nat. Rev. Genet. 21, 292–310 (2020).

[7] Lindblad-Toh, K. et al. A high-resolution map of human evolutionary constraint using 29 mammals. Nature 478, 476–482 (2011).

[8] Zoonomia Consortium. A comparative genomics multitool for scientific discovery and conservation. Nature 587, 240–245 (2020).

[9] Moore, J. E. et al. Expanded encyclopaedias of DNA elements in the human and mouse genomes. Nature 583, 699–710 (2020).

[10] Boix, C. A., James, B. T., Park, Y. P., Meuleman, W. & Kellis, M. Regulatory genomic circuitry of human disease loci by integrative epigenomics. Nature 590, 300–307 (2021).

[11] Claussnitzer, M. et al. A brief history of human disease genetics. Nature 577, 179–189 (2020).

[12] Madelaine, R. et al. A screen for deeply conserved non-coding GWAS SNPs uncovers a MIR-9-2 functional mutation associated to retinal vasculature defects in human. Nucleic Acids Res. 46, 3517–3531 (2018).

[13] Finucane, H. K. et al. Partitioning heritability by functional annotation using genome-wide association summary statistics. Nat. Genet. 47, 1228–1235 (2015).

[14] Kircher, M. et al. A general framework for estimating the relative pathogenicity of human genetic variants. Nat. Genet. 46, 310–315 (2014).

[15] Gulko, B., Hubisz, M. J., Gronau, I. & Siepel, A. A method for calculating probabilities of fitness consequences for point mutations across the human genome. Nat. Genet. 47, 276–283 (2015).

[16] Siepel, A. et al. Evolutionarily conserved elements in vertebrate, insect, worm, and yeast genomes. Genome Res. 15, 1034–1050 (2005).

[17] Short, P. J. et al. De novo mutations in regulatory elements in neurodevelopmental disorders. Nature 555, 611–616 (2018).

[18] Gjoneska, E. et al. Conserved epigenomic signals in mice and humans reveal immune basis of Alzheimer’s disease. Nature 518, 365–369 (2015).

[19] Hook, P. W. & McCallion, A. S. Leveraging mouse chromatin data for heritability enrichment informs common disease architecture and reveals cortical layer contributions to schizophrenia. Genome Res. 30, 528–539 (2020).

[20] Li, Y. E. et al. An atlas of gene regulatory elements in adult mouse cerebrum. Nature 598, 129–136 (2021).

[21] Srinivasan, C. et al. Addiction-associated genetic variants implicate brain cell type-and region-specific cis-regulatory elements in addiction neurobiology. J. Neurosci. 41, 9008–9030 (2021).

[22] Villar, D. et al. Enhancer evolution across 20 mammalian species. Cell 160, 554–566 (2015).

[23] Hujoel, M. L., Gazal, S., Hormozdiari, F., van de Geijn, B. & Price, A. L. Disease heritability enrichment of regulatory elements is concentrated in elements with ancient sequence age and conserved function across species. Am. J. Hum. Genet. 104, 611–624 (2019).

[24] Marnetto, D. et al. Evolutionary rewiring of human regulatory networks by waves of genome expansion. Am. J. Hum. Genet. 102, 207–218 (2018).

[25] Hardison, R. C., Oeltjen, J. & Miller, W. Long human–mouse sequence alignments reveal novel regulatory elements: A reason to sequence the mouse genome. Genome Res. 7, 959–966 (1997).

[26] Wasserman, W. W., Palumbo, M., Thompson, W., Fickett, J. W. & Lawrence, C. E. Human-mouse genome comparisons to locate regulatory sites. Nat. Genet. 26, 225–228 (2000).

[27] Lupo, G. et al. Molecular profiling of aged neural progenitors identifies Dbx2 as a candi-date regulator of age-associated neurogenic decline. Aging Cell 17, e12745 (2018).

[28] Davydov, E. V. et al. Identifying a high fraction of the human genome to be under selective constraint using GERP++. PLoS Comput. Biol. 6, e1001025 (2010).

[29] Amariuta, T. et al. Improving the trans-ancestry portability of polygenic risk scores by prioritizing variants in predicted cell-type-specific regulatory elements. Nat. Genet. 52, 1346–1354 (2020).

[30] Poch, T. et al. Single-cell atlas of hepatic T cells reveals expansion of liver-resident naivelike CD4+ T cells in primary sclerosing cholangitis. J. Hepatol. 75, 414–423 (2021).

[31] Yamanaka, Y. et al. Blocking fibrotic signaling in fibroblasts from patients with carpal tunnel syndrome. J. Cell. Physiol. 233, 2067–2074 (2018).

[32] Nasser, J. et al. Genome-wide enhancer maps link risk variants to disease genes. Nature 593, 238–243 (2021).

[33] Zhu, X., Duren, Z. & Wong, W. H. Modeling regulatory network topology improves genome-wide analyses of complex human traits. Nat. Commun. 12, 2851 (2021).

[34] Yengo, L. et al. Meta-analysis of genome-wide association studies for height and body mass index in 700000 individuals of European ancestry. Hum. Mol. Genet. 27, 3641–3649 (2018).

[35] Pardiñas, A. F. et al. Common schizophrenia alleles are enriched in mutation-intolerant genes and in regions under strong background selection. Nat. Genet. 50, 381–389 (2018).

[36] Zhu, X. & Stephens, M. Large-scale genome-wide enrichment analyses identify new trait-associated genes and pathways across 31 human phenotypes. Nat. Commun. 9, 4361 (2018).

[37] Loos, R. J. & Yeo, G. S. The genetics of obesity: from discovery to biology. Nat. Rev. Genet. 23, 120–133 (2022).

[38] Birnbaum, R. & Weinberger, D. R. Genetic insights into the neurodevelopmental origins of schizophrenia. Nat. Rev. Neurosci. 18, 727–740 (2017).

[39] Gulyaeva, O., Nguyen, H., Sambeat, A., Heydari, K. & Sul, H. S. Sox9-Meis1 inactivation is required for adipogenesis, advancing Pref-1+ to PDGFRα+ cells. Cell Rep. 25, 1002–1017 (2018).

[40] Owa, T. et al. Meis1 coordinates cerebellar granule cell development by regulating Pax6 transcription, BMP signaling and Atoh1 degradation. J. Neurosci. 38, 1277–1294 (2018).

[41] Bertrand, C., Valet, P. & Castan-Laurell, I. Apelin and energy metabolism. Front. Physiol. 6, 115 (2015).

[42] Castan-Laurell, I. et al. Apelin, diabetes, and obesity. Endocrine 40, 1–9 (2011).

[43] Beanan, M. J. & Sargent, T. D. Regulation and function of Dlx3 in vertebrate development. Dev. Dyn. 218, 545–553 (2000).

[44] Pao, P.-C. & Tsai, L.-H. Three decades of Cdk5. J. Biomed. Sci. 28, 79 (2021).

[45] Choi, J. H. et al. Anti-diabetic drugs inhibit obesity-linked phosphorylation of PPARγ by Cdk5. Nature 466, 451–456 (2010).

[46] Magen, D. et al. Autosomal recessive lissencephaly with cerebellar hypoplasia is associated with a loss-of-function mutation in CDK5. Hum. Genet. 134, 305–314 (2015).

[47] Pereira, C., Azevedo, I., Monteiro, R. & Martins, M. 11β-Hydroxysteroid dehydrogenase type 1: relevance of its modulation in the pathophysiology of obesity, the metabolic syndrome and type 2 diabetes mellitus. Diabetes Obes. Metab. 14, 869–881 (2012).

[48] Masuzaki, H. et al. A transgenic model of visceral obesity and the metabolic syndrome. Science 294, 2166–2170 (2001).

[49] Lawson, A. J. et al. Cortisone-reductase deficiency associated with heterozygous mutations in 11β-hydroxysteroid dehydrogenase type 1. Proc. Natl. Acad. Sci. U.S.A. 108, 4111–4116 (2011).

[50] Akalestou, E., Genser, L. & Rutter, G. A. Glucocorticoid metabolism in obesity and following weight loss. Front. Endocrinol. 11, 59 (2020).

[51] McEvilly, R. J., de Diaz, M. O., Schonemann, M. D., Hooshmand, F. & Rosenfeld, M. G. Transcriptional regulation of cortical neuron migration by POU domain factors. Science 295, 1528–1532 (2002).

[52] Kuwahara, A. et al. Tcf3 represses Wnt-β-catenin signaling and maintains neural stem cell population during neocortical development. PLoS ONE 9, e94408 (2014).

[53] Riley, P., Anaon-Cartwight, L. & Cross, J. C. The Hand1 bHLH transcription factor is essential for placentation and cardiac morphogenesis. Nat. Genet. 18, 271–275 (1998).

[54] Ursini, G. et al. Convergence of placenta biology and genetic risk for schizophrenia. Nat. Med. 24, 792–801 (2018).

[55] Khandaker, G. M. et al. Inflammation and immunity in schizophrenia: implications for pathophysiology and treatment. Lancet Psychiatry 2, 258–270 (2015).

[56] Splawski, I. et al. CaV1.2 calcium channel dysfunction causes a multisystem disorder including arrhythmia and autism. Cell 119, 19–31 (2004).

[57] Chen, C. et al. Ciliopathy protein HYLS1 coordinates the biogenesis and signaling of primary cilia by activating the ciliary lipid kinase PIPKIγ. Sci. Adv. 7, eabe3401 (2021).

[58] Liu, S., Trupiano, M. X., Simon, J., Guo, J. & Anton, E. The essential role of primary cilia in cerebral cortical development and disorders. Curr. Top. Dev. Biol. 142, 99–146 (2021).

[59] Djenoune, L., Berg, K., Brueckner, M. & Yuan, S. A change of heart: new roles for cilia in cardiac development and disease. Nat. Rev. Cardiol. 19, 211–227 (2022).

[60] Mee, L. et al. Hydrolethalus syndrome is caused by a missense mutation in a novel gene HYLS1. Hum. Mol. Genet. 14, 1475–1488 (2005).

[61] Matthijs, G. et al. Mutations in PMM2, a phosphomannomutase gene on chromosome 16p13 in carbohydrate-deficient glycoprotein type I syndrome (Jaeken syndrome). Nat. Genet. 16, 88–92 (1997).

[62] Grünewald, S. The clinical spectrum of phosphomannomutase 2 deficiency (CDG-Ia). Biochim. Biophys. Acta - Mol. Basis Dis. 1792, 827–834 (2009).

[63] Loaeza-Reyes, K. J. et al. An overview of glycosylation and its impact on cardiovascular health and disease. Front. Mol. Biosci. 8, 751637 (2021).

[64] Rebelo, A. L., Chevalier, M. T., Russo, L. & Pandit, A. Role and therapeutic implications of protein glycosylation in neuroinflammation. Trends Mol. Med. 28, 270–289 (2022).

[65] Williams, S. E., Mealer, R. G., Scolnick, E. M., Smoller, J. W. & Cummings, R. D. Aberrant glycosylation in schizophrenia: a review of 25 years of post-mortem brain studies. Mol. Psychiatry 25, 3198–3207 (2020).

[66] Correll, C. U. et al. Prevalence, incidence and mortality from cardiovascular disease in patients with pooled and specific severe mental illness: a large-scale meta-analysis of 3,211,768 patients and 113,383,368 controls. World Psychiatry 16, 163–180 (2017).

[67] Snetkova, V. et al. Ultraconserved enhancer function does not require perfect sequence conservation. Nat. Genet. 53, 521–528 (2021).

[68] Wong, E. S. et al. Deep conservation of the enhancer regulatory code in animals. Science 370, eaax8137 (2020).

[69] Kwon, S. B. & Ernst, J. Learning a genome-wide score of human-mouse conservation at the functional genomics level. Nat. Commun. 12, 2495 (2021).

[70] Zhang, K. et al. A single-cell atlas of chromatin accessibility in the human genome. Cell 184, 5985–6001 (2021).

[71] Bartosovic, M., Kabbe, M. & Castelo-Branco, G. Single-cell CUT&Tag profiles histone modifications and transcription factors in complex tissues. Nat. Biotechnol. 39, 825–835 (2021).

[72] Ma, S., Chen, X., Zhu, X., Tsao, P. S. & Wong, W. H. Leveraging cell-type-specific regulatory networks to interpret genetic variants in abdominal aortic aneurysm. Proc. Natl. Acad. Sci. U.S.A. 119, e2115601119 (2022).

[73] Ramdas, S. et al. A multi-layer functional genomic analysis to understand noncoding genetic variation in lipids. Am. J. Hum. Genet. 109, 1366–1387 (2022).

[74] Lin, X. et al. Nested epistasis enhancer networks for robust genome regulation. Science https://doi.org/10.1126/science.abk3512 (2022).

[75] Kuhn, R. M., Haussler, D. & Kent, W. J. The UCSC genome browser and associated tools. Brief. Bioinform. 14, 144–161 (2013).

[76] Quinlan, A. R. & Hall, I. M. BEDTools: a flexible suite of utilities for comparing genomic features. Bioinformatics 26, 841–842 (2010).

[77] 1000 Genomes Project Consortium. A global reference for human genetic variation. Nature 526, 68–74 (2015).

[78] Benner, C. et al. FINEMAP: efficient variable selection using summary data from genome-wide association studies. Bioinformatics 32, 1493–1501 (2016).

[79] Wang, G., Sarkar, A., Carbonetto, P. & Stephens, M. A simple new approach to variable selection in regression, with application to genetic fine mapping. J. R. Stat. Soc., B: Stat. Methodol. 82, 1273–1300 (2020).

[80] Zhu, X. & Stephens, M. Bayesian large-scale multiple regression with summary statistics from genome-wide association studies. Ann. Appl. Stat. 11, 1561 (2017).

[81] Heinz, S. et al. Simple combinations of lineage-determining transcription factors prime cis-regulatory elements required for macrophage and B cell identities. Mol. Cell 38, 576–589 (2010).

[82] Duren, Z., Chen, X., Jiang, R., Wang, Y. & Wong, W. H. Modeling gene regulation from paired expression and chromatin accessibility data. Proc. Natl. Acad. Sci. U.S.A. 114, E4914–E4923 (2017).

[83] Yousefi, S. et al. Comprehensive multi-omics integration identifies differentially active enhancers during human brain development with clinical relevance. Genome Med. 13, 162 (2021).

[84] Wang, D. et al. Comprehensive functional genomic resource and integrative model for the human brain. Science 362, eaat8464 (2018).

[85] Bult, C. J., Blake, J. A., Smith, C. L., Kadin, J. A. & Richardson, J. E. Mouse genome database (MGD) 2019. Nucleic Acids Res. 47, D801–D806 (2019).

[86] Zhou, Y. et al. Metascape provides a biologist-oriented resource for the analysis of systems-level datasets. Nat. Commun. 10, 1523 (2019).

[87] Amberger, J. S., Bocchini, C. A., Scott, A. F. & Hamosh, A. OMIM.org: leveraging knowledge across phenotype-gene relationships. Nucleic Acids Res. 47, D1038–D1043 (2019).

[88] Zhou, Y. et al. Therapeutic target database update 2022: facilitating drug discovery with enriched comparative data of targeted agents. Nucleic Acids Res. 50, D1398–D1407 (2022).

[89] Buniello, A. et al. The NHGRI-EBI GWAS Catalog of published genome-wide association studies, targeted arrays and summary statistics 2019. Nucleic Acids Res. 47, D1005–D1012 (2019).

